# Chromosomal breakage in *Arabidopsis thaliana* neo-polyploids occurs preferentially at candidate fragile sites colocalized with ancient genome rearrangements

**DOI:** 10.64898/2026.09.21.753295

**Authors:** Thanvi Srikant, Hui San Tan, Kirsten Bomblies, Adrián Gonzalo

## Abstract

Newly formed polyploids often suffer extensive aneuploidy and chromosome rearrangement. We explored whether these are predictable and conserved across genotypes. We quantified whole-chromosome and segmental aneuploidy in progeny arrays of neo-polyploid *Arabidopsis thaliana* from two accessions (Col-0 and L*er*). 29% of progeny had whole-chromosome and/or segmental aneuploidies. Strikingly, segmental aneuploidy arose exclusively from L*er*, preferentially maternally, showing that genotype and sex of origin matter. Breakpoints of segmental aneuploidies were non-random, with over 2/3 mapping to five potentially fragile regions (PFRs). Two PFRs overlap with junctions of ancient chromosomal fusions that differentiate the *A. thaliana* genome from that of its relatives and also overlap more broadly with hotspots of recurrent chromosomal rearrangements differentiating species in the *Brassicaceae* family. Several PFRs overlap with large tandemly repeated disease resistance gene clusters. Our results support the hypothesis of a link between ancestral genome rearrangements and present-day chromosome fragility. Polyploidy serving as a context that makes aneuploidies more likely to arise and survive, with fragile sites lending a degree of predictability.

**GRAPHICAL ABSTRACT:** 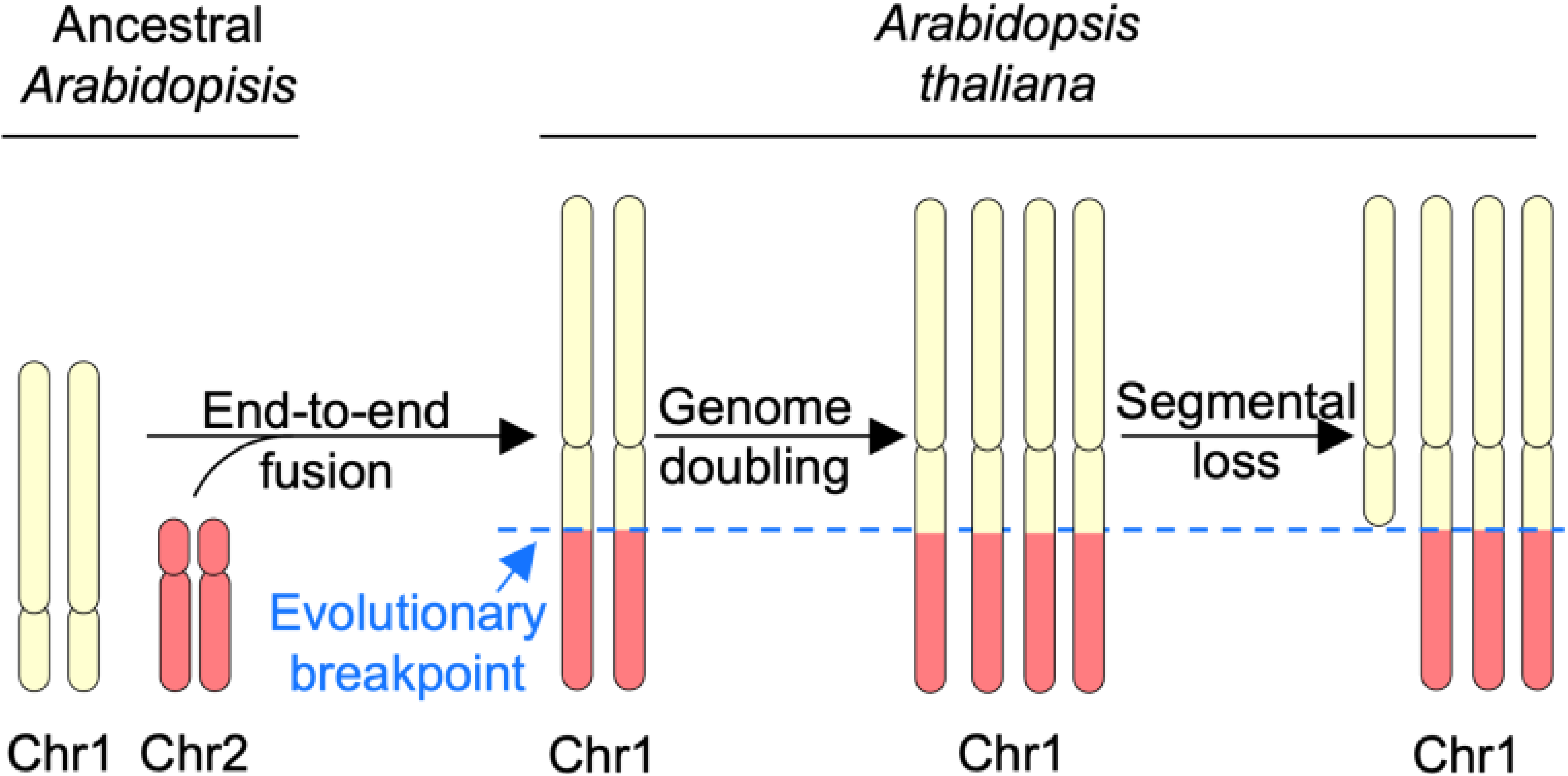

**HIGHLIGHTS:**

- 29% of neo-polyploid *A. thaliana* progeny are chromosomal or segmental aneuploids.
- Comparing two strains, segmental aneuploidies originated only from one parent.
- Breakpoints cluster in fragile regions (PFRs) overlapping ancient rearrangements.
- Three of five PFRs colocalize with large disease resistance gene clusters.

## INTRODUCTION

Polyploidy is thought to be a major driver of adaptive innovation, which might explain its pervasiveness in evolution and in crops ^1–3^. However, doubling the chromosome complement often causes problems with genome stability ^4–12^. Newly formed autopolyploids commonly have an increased propensity to chromosome mis-segregation and breakage in meiosis, but are generally more tolerant of altered dosage than diploids, making it more likely that these events are observed in progeny ^12–16^. While aneuploidy is generally deleterious ^17,18^, there is also evidence that it can occasionally be a source of innovation and evolvability ^17,19^.

Polyploids tend to undergo extensive segmental losses and other rearrangements, which in the longer term can lead to re-diploidization of the genome ^20–25^. Whole genome duplication thus serves as an important source of structural variation in genomes ^5,11,12,16,19,26^. It is not yet clear whether these deletions arise randomly, or whether some regions of the genome are particularly prone to them. In animals, where the causes of karyotypic variation have been extensively studied, breakpoints of rearrangements that differ among species often overlap fragile sites that are prone to chromosome breakage within species ^27,27,28,28–32^. Thus, the fragility of certain regions has been suggested as an explanation for why certain breakpoints are often reused in independent genome rearrangements across animal diversity ^28,33,34^. Something similar may be going on in plants, though the evidence to date is much less extensive. For example, breakpoint reuse has been documented in a mutagenesis study in *A. thaliana* where a common deletion breakpoint region corresponded to a breakpoint of interspecies rearrangements ^35^. In wheat, breakpoints of an ancestral inversion are reused in rearrangements differentiating later-arising species ^36^, while in another grass, *Phleum echinatum*, repetitive regions and intrachromosomal telomere repeats appear to be fragile, potentially mapping to junctions of karyotypic differences among species ^37^.

Polyploids, due to their increased genomic instability as well as their greater tolerance to aneuploidy, provide a useful system to explore whether loss/gain patterns of genetic material are random or not. We sought to better understand the patterns of heritable post-polyploidization aneuploidy in neo-tetraploid *A. thaliana* of two commonly used genetically distinct accessions (Col-0 and L*er*). This allowed us to ask whether there are genotypic or sex differences in the types or rates of both full-length and/or segmental aneuploidies. Consistent with previous studies ^4^, we find whole-chromosome aneuploidy affects over 25% of progeny, with non-significant differences by genotype or sex. We also found segmental deletions and duplications, but these originated exclusively from the L*er* parent, more often maternally. Around 2/3 of segmental aneuploidy breakpoints were clustered into five potentially fragile regions (PFRs). Interestingly, two of these PFRs co-localise to breakpoint regions of genomic rearrangements that occurred during the evolution of *A. thaliana* and in some cases, the Brassicaceae family more broadly. Thus, it seems that plants, as has been reported in animals ^27,28,32^, have multiple fragile chromosome regions in their genomes that are not only prone to breakage within species in aberrant situations like neo-polyploidy or replication stress, but can also contribute to karyotypic evolution among species.

## RESULTS

### Aneuploidy in the progeny of neo-tetraploid *Arabidopsis thaliana*

We characterised the types and quantity of aneuploid progeny produced by neo-tetraploid *Arabidopsis thaliana* by analysing progeny arrays from reciprocal crosses between neo-tetraploids of the Col-0 and L*er* accessions following a previously developed read coverage-based approach ^38,39^. We generated tetraploids using mutants that yield unreduced gametes (*ps1, tam1, osd1*, see Figure S1A) to avoid the use of colchicine, which usually produces chimeric mixed-ploidy plants. We reciprocally crossed the resulting L*er* and Col-0 neo-tetraploid plants (*A. thaliana* strains are highly homozygous, so gametes are genetically uniform in terms of genome sequence) to generate an F_1_ population of 228 individuals (Figure 1A, Figure S1A), of which 131 had a Col-0 mother and a L*er* father, and 97 the reverse. We sequenced all 228 F_1_ individuals using short-read sequencing. We included diploid and tetraploid inbreds (Figure S1) and three diploid Col-0 x Ler F_1_ individuals to allow us to correct for differences in read coverage across the genome.

**Figure 1.**
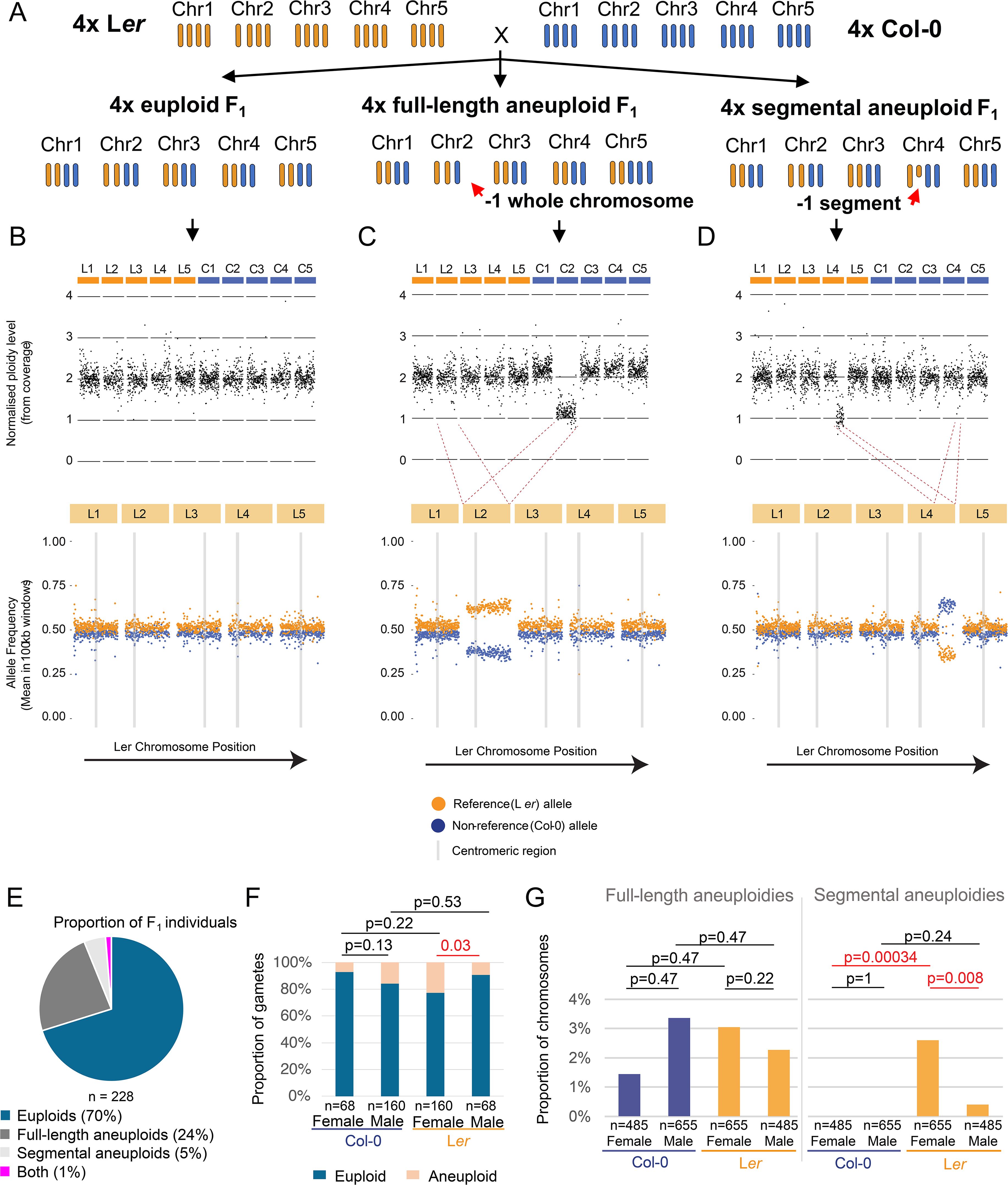
Detection and analyses of aneuploidies in neo-tetraploid *A. thaliana*. (A) Schematic representation of how neo-tetraploid F_1_ were generated through reciprocal crossing of Col-0 and L*er* neo-tetraploid inbreds. See Figure S1A for more details. (B-D) Examples of aneuploidy detection using sequencing coverage data from three F_1_ individuals, “Cross_3_Plant_32”, “Cross_3_Plant_25” and “Cross_38_Plant_01”, respectively (Figure S2, S3). The upper panels show aneuploidy detection using a read-coverage approach and the lower panels show a variant-calling based approach. The Ler chromosomes are indicated as L1-L5 (orange) and Col-0 chromosomes are indicated as C1-C5 (blue). In both panels, every data point represents a mean value (normalised read coverage / allele frequency) in a 100kb window of the genome. For the upper panels, sequencing data are mapped to an in-silico hybrid genome that combines TAIR10.1 (Col-0) and Ler. For the lower panels, the sequencing data are mapped to a Ler reference genome. (B) Data from a euploid individual with normalised coverage consistent with two copies of each maternally and paternally inherited chromosome (upper panel), and allele frequencies clustering around 0.5 when mapped to a Ler reference genome (lower panel). (C) Data from an individual affected by a full-length aneuploidy – a loss of one Chromosome 2 copy from Col-0, resulting in a drop in coverage (upper panel) and sharp changes in allele frequency (lower chromosome) for Chr2. (D) Data from an individual affected by a segmental loss of part of Chromosome 4 from the L*er* parent. (E) Pie chart showing the proportion of sequenced F_1_ individuals that are euploid, whole-chromosome aneuploid, segmental aneuploid, or both. (F) Bar graph showing the proportion of euploid and aneuploid inferred gametes that formed the studied F_1_ individuals (n = number of gametes in each class). (G) shows the proportion of chromosomes analysed that were affected by full-length or segmental aneuploidies separately based on their parent-of-origin (n = number of chromosomes in each class). For panels F and G, p-values above horizontal bars are from Fisher’s Exact tests for a given comparison. Note that repeated (identical) p-values are an artifact of False Discovery Rate correction.

To detect aneuploidies, we followed established approaches for molecular karyotyping using normalized sequencing read coverage (Figure S2) and allele frequencies (Figure S3) ^38,39^ to detect the presence, type, location, and parent of origin of aneuploidies (Figure 1A-D, Supplemental Data S1, see Methods). We first confirmed that the neo-tetraploid parents were indeed fully euploid (Figure S1B-C) and then proceeded to analyse the F_1_ progeny, quantifying both the proportions of aneuploid individuals and individual chromosomes affected by aneuploidy (note that the aneuploid individuals we identified can in principle have more than one aneuploid chromosome).

Among the 228 sequenced F_1_ individuals, 71% were tetraploid euploids (without detected aneuploidies) and 29% were aneuploids, which aligned well with numbers obtained from previous studies ^4^. 24% of individuals were affected by full-length aneuploidy, 5% by segmental aneuploidy, and 1% by both. The latter is what we expect if segmental and whole chromosome aneuploidy events occur independently of each other (Figure 1E).

We used a sequencing coverage-based aneuploidy detection approach differentiating Col-0 and Ler chromosomes of the 228 sequenced F_1_ individuals (2280 chromosomes analysed in total since each individual normally has 10 chromosomes, Figures S2-S3, Table S1). We identified 79 chromosomes affected by aneuploidy (hereafter referred to as “aneuploid chromosomes”), of which 60 were gains or losses of an entire chromosome (i.e. full-length aneuploidies; 76% of the aneuploid chromosomes). We also detected 19 chromosomes with segmental aneuploidy (24% of aneuploid chromosomes) involving gains or losses of parts of a chromosome.

To study the origin of aneuploidy, we inferred the constitution of the gametes that gave rise to each F_1_ individual (Supplemental Table S2). In terms of sex differences, in the L*er* accession, aneuploid gametes were significantly more frequent from maternal than paternal origin (23% vs 9% of inferred gametes, p = 0.03, Fisher’s exact test, Figure 1F), while Col-0 showed a weak non-significant trend in the opposite direction (p = 0.13, Figure 1F). Overall, there was no significant difference between accessions in the proportion of inferred aneuploid gametes (p > 0.22, Fisher’s exact test, Figure 1F). When considering individual chromosomes, we found that among the 79 aneuploidies identified (Supplemental Tables S3-S5), although there were differences among chromosomes in aneuploidy rate, the differences were not significant (p > 0.06, Fisher’s exact test, Figure S4A), perhaps due to the low frequency of aneuploidy events for any given chromosome. We did observe a preference for gains over losses for full-chromosome aneuploidies (67% gains, p = 0.013, Fisher’s exact test, Supplemental Tables S6, S7), mirroring trends reported in other studies^4,40^. This preference was evident regardless of sex or genotype of the parent (p = 0.17, Fisher’s exact test, Figure S4B). Since gains and losses due to chromosome mis-segregation should in principle be equally frequent, this difference could imply there is stronger selection against losses during the haploid gametophyte stage or early embryo development. Perhaps speaking to the former, from the female side, gains and losses are closer to the expected 50% than on the male side, suggesting there may be stronger selection against losses on the male side (Figure S4B).

In contrast to full-chromosome aneuploidies, for segmental aneuploidies there was a dramatic difference between accessions (Supplemental Tables S5, S8, S9); 100% of the 19 detected segmental aneuploidies originated from the L*er* parent (Figure 1G) and showed a significant bias for being of maternal origin (p = 0.008, Figure 1G). Most (16 of 24) involved losses from a breakpoint to the chromosome end, while the rest were interstitial (Figure 2A-C). In contrast to whole chromosome aneuploidy, losses were more common than gains (18 losses, 6 gains), though this is not true of the interstitial events (5 gains, 4 losses; Figure 2C). In rare cases, we found evidence for complex rearrangements with multiple adjacent changes in copy number (e.g. Figure S5A-D). Note that these rates of segmental aneuploidies are conservative estimates of the rates of rearrangement since our experimental design does not allow us to identify inversions or translocations that do not alter copy number.

**Figure 2.**
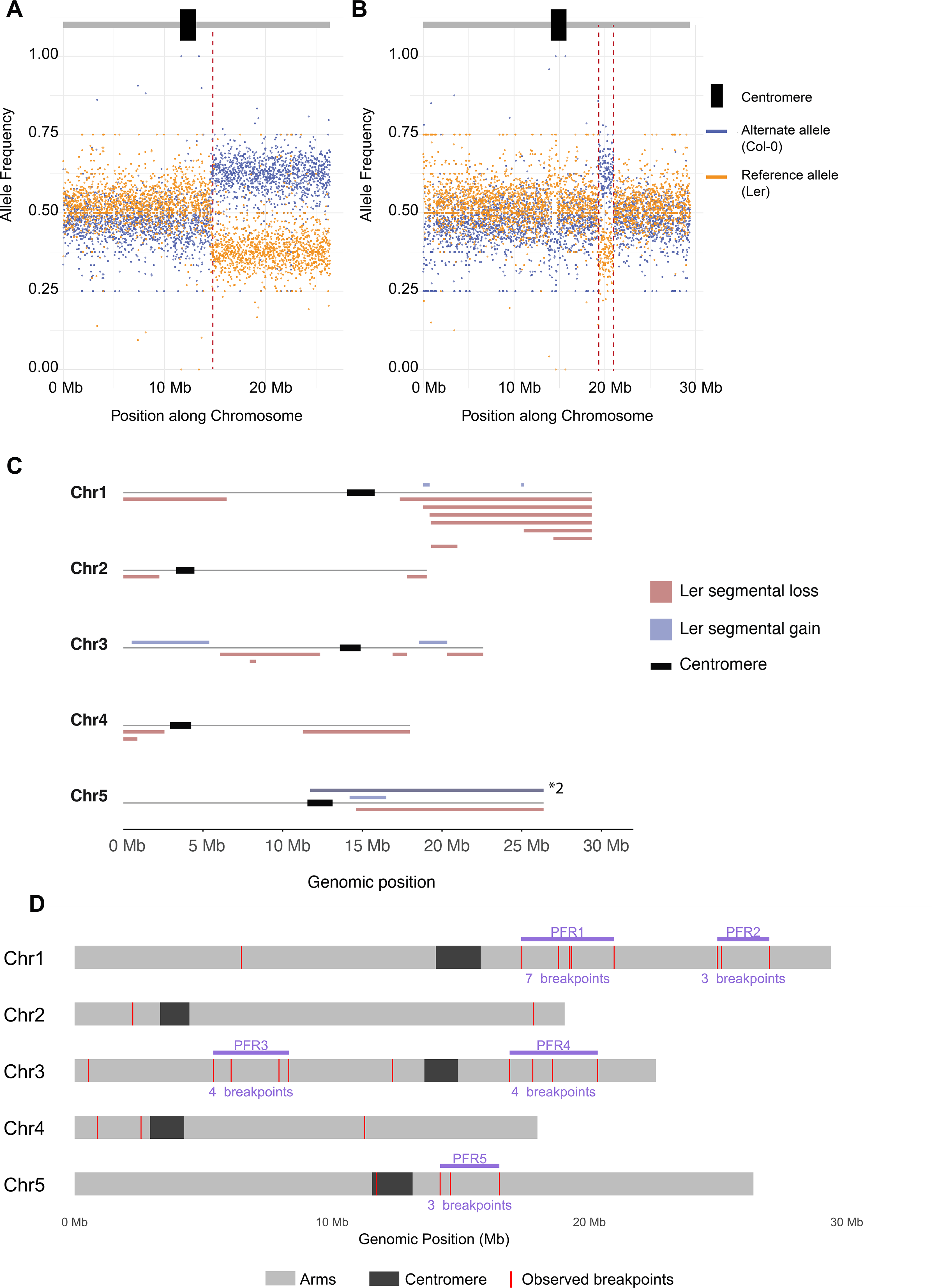
Breakpoints in segmental aneuploids. (A, B) Two examples of how breakpoints are mapped using allele frequencies at high resolution mapped to the Ler reference genome. Each data point represents the average value for a 10 kb window across the chromosome. Panel A shows Chr1 from individual “Cross_6_Plant_29” and panel B shows Chr5 from individual “Cross_4_Plant_45” (Figure S2, S3). (C) Shows the size and position of all segmental gains and losses we identified. The *2 label indicates that the same segmental gain was observed in double dosage in one individual. (D) A breakpoint map of L*er* genome, showing the identified breakpoints as red lines, and PFRs as purple lines, with the number of breakpoints in each shown below.

### Potentially fragile regions (PFRs) in *A. thaliana* chromosomes

We next examined the distribution of the segmental aneuploidy breakpoints. In the 19 chromosomes affected by segmental aneuploidy, we identified 18 segmental losses and 6 gains, with some affected chromosomes carrying two or more different rearrangements. Based on changes in allele frequencies, we mapped 30 breakpoints at a resolution of 10 kb (Figure 2A-C, Supplemental Table S10). Nearly all of them mapped with distinct coordinates at fine scale, but two breakpoints in different plants from the same cross mapped to the same 10 kb window. This opens the possibility that some breakpoints might reflect somatic events inherited to different individuals. This analysis showed that the size of the segmental aneuploidies (Figure 2C, Supplemental Table S11) ranged from 160 kb (Figure S5C-D) to 14.6 Mb for segmental gains and from 360 kb to 12 Mb for segmental losses.

We next visualized the locations of all 30 mapped breakpoints (Figure 2D, Supplemental Table S11). We noticed that one 3.7 Mb region of Chromosome 1 contained seven breakpoints corresponding to six independent segmental losses or gains (Figure 2C, D). This disproportionate concentration of breakpoints in a relatively small fraction of the genome suggests that this genomic region might be fragile, i.e. more likely to undergo a chromosomal break than the genome average. To ask if there might be other potentially fragile regions of similar size, we searched for additional regions up to 3.7 Mb in length with at least three breakpoints (which would be significantly more than expected by chance; p = 0.01, binomial test). Since the windows were not chosen independently of breakpoint positions (i.e. chosen *a posteriori*), statistical tests are not valid to conclude whether these windows are indeed enriched in breakpoints. Therefore, we cautiously named these as ‘Potentially Fragile Regions’ (PFRs). We identified five such regions (Figure 2D, Supplemental Table S12); which harboured seven (the aforementioned PFR1), three (PFR2), four (PFR3), four (PFR4) and three (PFR5) breakpoints, respectively. Thus, 21 of the 30 detected breakpoints fell within five PFR regions.

### PFRs associate with recurrent historic chromosome rearrangements

The genome structure of *A. thaliana* differs from that of its relatives in multiple end-to-end chromosome fusions, inversions, and other rearrangements ^21,41–44^ (Figure 3A). We observed that the genomic location of PFR1, which had the highest number of independent breakpoints, positionally overlaps with the junction of a known ancient end-to-end fusion (‘EF’) between two ancestral chromosomes (Chromosomes 1 and 2 in related lineages; Figure 3A and B) that took place less than 12 MYA, after the divergence of *A. thaliana* and the rest of the *Arabidopsis* species ^21,43,45–47^. To ask how concomitant our *de novo* breakpoints and the ancient breakpoint really are, we mapped the precise coordinates of the EF using microsynteny with related species, all of which retain the ancestral arrangement (*Arabidopsis halleri*, *Arabidopsis lyrata*, *Arabidopsis arenosa*, *Capsella rubella* and *Cardamine amara*). We observed a complex array of nested rearrangements among species at the EF site (Figure 3A, B; Figure S6). Strikingly, the EF site is located exactly within the core of PFR1, the central 0.5 Mb region where five out of the seven PFR1 breakpoints are concentrated (Figure 3B, Figure S6). PFR5 also co-localizes with an EF event (Figure 3A). This suggests that boundaries of at least some of the ancestral rearrangement events remain prone to breakage millions of years later.

**Figure 3.**
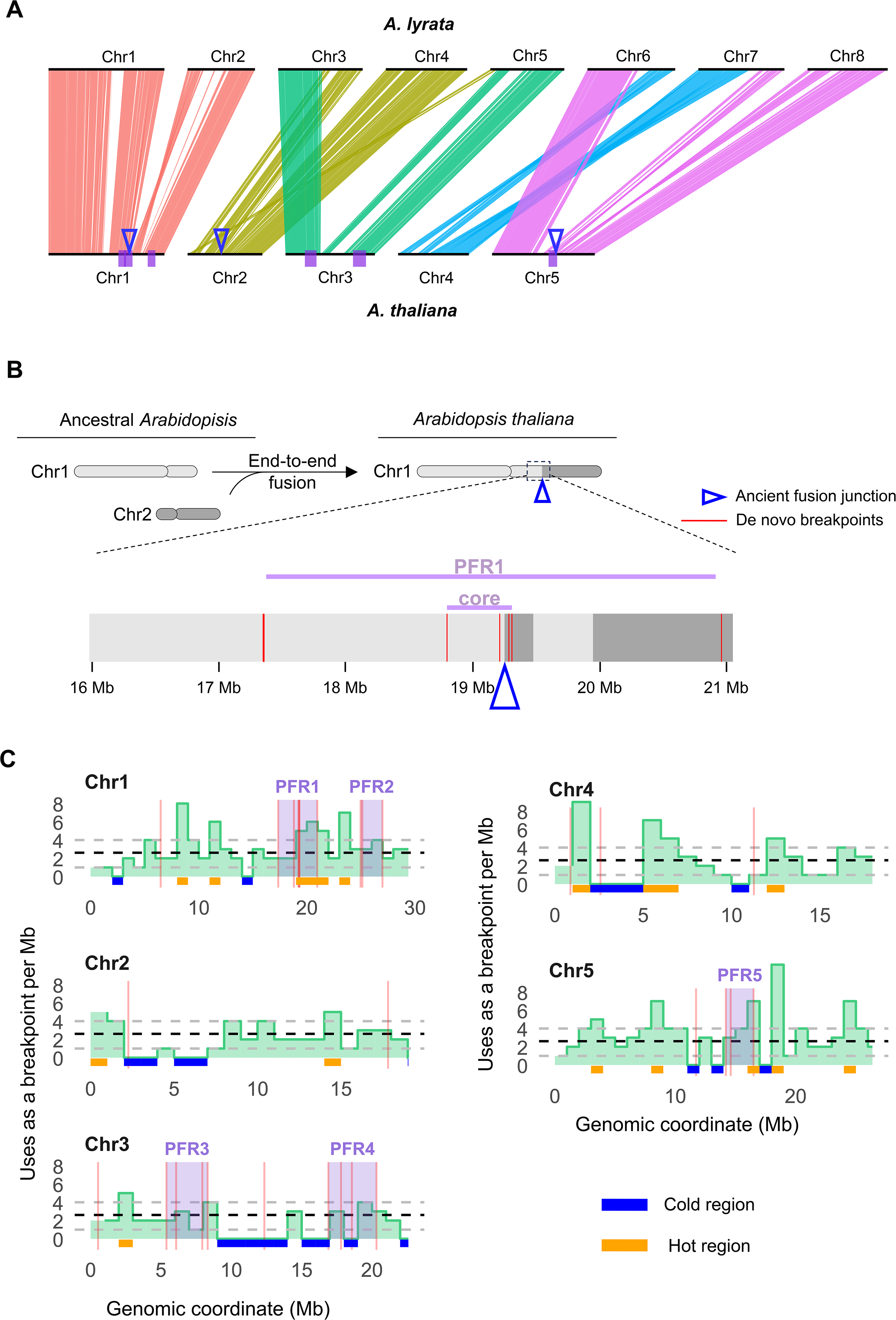
Links between PFRs and ancient rearrangements in the Brassicaceae family. (A) Synteny plot of the five chromosomes of *A. thaliana* and the eight in *A. lyrata* (redrawn from data in ^85^). Triangles indicate the position of the end-to-end fusion junctions on *A. thaliana* chromosomes. PFRs are highlighted in purple under *A. thaliana* chromosomes. (B) Close-up of the mapped breakpoints in PFR1 (vertical red lines) and its core region, overlapping with the end-to-end fusion junction in *A. thaliana* Chromosome 1, color-coded to show chromosome origin; dark and light gray indicate sequence origins from ancestral chromosomes 1 and 2, respectively. (C) Density map of the breakpoints for ancient genome rearrangements identified during 25 MY of *Brassicaceae* evolution in ^43^. The genome is divided in bins of 1 Mb. Black dashed lines indicate the genome average of ancient breakpoints per Mb whereas the lower and upper grey dashed lines indicate the first and the fourth quantiles (respectively) for ancient breakpoints per megabase.

All 11 breakpoints we identified in Chromosome 1 were significantly nearer to the ancient EF site than expected by chance (p = 0.0008, permutation test, Figure S7). We documented a similar trend for the EF located on Chromosome 5 (p = 0.0081, permutation test, Figure S7), but not for the one on Chromosome 2 (p = 0.54, permutation test, Figure S7). The observation that two of the three EFs in *A. thaliana* coincide with a PFR is consistent with animal studies where evolutionary breakpoint regions often coincide with known intra-species fragile sites ^27,28,48^.

We also asked whether the evolutionary breakpoints of even more ancient rearrangements that differentiate species within the Brassicaceae more broadly could also be associated with PFRs in *A. thaliana*, as has been observed in animals ^27,28,32,49^. We therefore referred to a recent, detailed reconstruction of rearrangements between synteny blocks spanning 25 million years of evolution in the *Brassicaceae* family ^43^. To enable the phylogenetic study of rearrangements, instead of precise breakpoints (which vary across species) consensus block boundaries are used to map approximate breakpoints of each block ^43^.

We counted how often each block boundary was involved in a historical rearrangement (see Methods; Figure 3C). We detected 15 ‘hot’ regions (continuous regions within the fourth quartile of the “reuse rate per Mb”; i.e. more than 4 uses per Mb) and 12 ‘cold’ regions (continuous regions within the first quartile; i.e. zero uses per Mb) for ancient rearrangements (Figure 3C; Supplemental Table S15). PFR1, the most fragile region we identified, has been used as an evolutionary breakpoint 15 times in the last 25 MYA and overlaps with a 3 Mb hot region that is the largest in length and has the greatest number (16) of total uses as an evolutionary breakpoint in the sampled species ^43^ (Figure 3C, Supplemental Table S15). In addition, PFR5, also overlaps with a 1 Mb hot region that was used 14 times as an evolutionary breakpoint in the sampled species (Figure 3C, Supplemental table S15). Nevertheless, though generally associated more with hot regions, some *de novo* breakpoints also took place within cold regions. Overall, this association between *de novo* and evolutionary breakpoints suggests that certain regions prone to be involved in genome rearrangements within species could remain fragile for extended periods of evolutionary time, suggesting their fragility is either unavoidable or functionally relevant as has been argued for animals ^52–54^.

### Genomic and epigenetic features of PFRs

Since multiple chromosome breakages occurred in regions involved in ancient genome rearrangements, we asked whether PFRs might share specific characteristics with known patterns at intraspecific fragile sites ^31^ or evolutionary breakpoint regions in animals ^27,28,32^. We assessed enrichment relative to genome-wide levels in gene density, AT nucleotide content, transposable elements (TEs), and chromatin accessibility (the latter using published ATAC-seq data from Ler wildtype plants ^55^ (Supplemental Table S18)). Comparing all the *de novo* breakpoints (using windows spanning 250 kb upstream and downstream of every mapped breakpoint; Supplemental Table S18), we found a significant depletion of genes in the vicinity of breakpoints (p = 0.0008, permutation test). This is in contrast to evolutionary breakpoint regions in animals, which are often gene-rich, but in agreement with patterns around many fragile sites in humans, which are typically gene-depleted ^27,28,32^. We found no significant trends for the other features we examined (p > 0.22, permutation test).

We also analysed just PFR regions and found that only PFR3 and PFR5 diverged significantly from the genome average for any of the tested genomic features. PFR3 showed higher chromatin accessibility in L*er* than the genome average (p = 0.0008, permutation test), while PFR5 had higher AT content (p = 0.046), which has also been observed at many fragile sites in human chromosomes ^31^ and many evolutionary breakpoint regions in animals ^27,28,32^. We also performed the same analysis at the core of PFR1 (Figure 2C, 3B), but no significant divergence from genome averages for any of the tested features was observed (p > 0.12, permutation test). In summary, these analyses showed that although individual PFRs might share some genomic or epigenetic features with evolutionary breakpoint regions and fragile sites in animals, there is no systematic trend for the features we tested for. This might suggest either that these PFRs may share some feature we did not test for, or that each PFR might be fragile for a different reason.

### Genes overrepresented at hot regions and PFRs

Given that genes within dynamic rearranged regions have been proposed to be associated with adaptation ^8,33,51,56–60^, we asked whether genes in regions that are highly rearranged during Brassicaceae karyotypic evolution were enriched for any biological processes. We found that gene ontology (GO) categories associated with response to environmental stresses are significantly enriched at hot regions. These categories include defense response, response to stimulus, response to abiotic stress, and regulation of biosynthetic processes, as well as other categories like recognition of pollen, pollen-pistil interaction and developmental processes involved in reproduction (Supplemental Table S16).

The defense-related GO category was particularly striking, since in plants many pathogen response genes of the nucleotide-binding and leucine-rich repeat (NLR) class are often organized as clusters of tandemly duplicated genes that have been associated with frequent rearrangements ^8,59–61^. NLR clusters are particularly abundant in *A. thaliana* on Chromosome 1 and Chromosome 5 ^59,62^, PFR1, PFR2 (both on Chromosome 1) and PFR5 (on Chromosome 5) were significantly enriched in NLR genes (p < 0.001, binomial one-tailed test), suggesting a potential association between fragility and clusters of NLR genes at these PFRs (Figure S8). Whether the dynamism of these NLR clusters is a cause or consequence of fragility in these regions will be an interesting question for future work.

## DISCUSION

In this study, we analyzed patterns of whole chromosome and segmental gains and losses in *Arabidopsis thaliana* neo-tetraploids of two accessions. The capacity of polyploids to buffer dosage changes provides a context in which such events are more likely to be inherited and thus detected in progeny arrays. Our reciprocal cross experimental design allowed us to track whether aneuploidy rates differ by sex or genotype of the parent of origin. The relatively high frequency of whole chromosome aneuploids we identified is similar to previous studies, and is likely attributable to meiotic instability leading to chromosome segregation errors, a well-known issue in neo-polyploid meiosis ^4,12,12,14,15^.

For whole-chromosome aneuploidy rates, we saw no significant difference between Col-0 and L*er* and rates or female vs. male, suggesting there is no consistent polyploidy-caused difference. Gains of chromosomes were more common than losses, especially from the male side, suggesting that perhaps selection at the level of gametophyte function works more strongly against chromosome loss. Although in previous work in *A. thaliana* neo-tetraploids, a similar bias for gain was observed ^4^, it did not seem to be more pronounced in male gametes. The trend we observed also contrasts results from *A. thaliana* diploids, where trisomies were found to be preferentially of maternal origin ^39^. One potential explanation for this latter result could be related to the “triploid block” effect, where fertility is especially strongly impaired by chromosomes of paternal origin ^63^, which may be less relevant in a polyploid context simply because there are more copies of each chromosome.

The segmental aneuploidies we detected could reflect chromosomal instability and breakage due to unrepaired breaks and entanglements ^12–16^. In our experiment, segmental aneuploidies were rarer than whole-chromosome aneuploidies, and remarkably, all originated from L*er*, with a strong bias toward female origin, and losses being more common than gains (except for interstitial events). Other examples of natural variation for susceptibility to chromosome breakage include maize strains with differences in transposon activity ^64^ and a recent GWAS in *A. thaliana* diploid accessions for genome content variability^65^. The accession-level difference could potentially be explained by L*er* having a greater susceptibility to breakage or a lesser ability to repair breaks. Indeed, the L*er* accession was obtained from irradiated seeds of a natural population of *A. thaliana* ^66^ and is known to carry a loss-of-function mutation in one of two copies of a key DNA repair helicase, *RECQ4B*, while both copies are active in Col-0 ^67^. Whether this mutation is causal in this case is not known, but our results underscore that genetic background can cause differences in the probability of segmental aneuploidies affecting progeny.

The most intriguing feature of the segmental aneuploidies is the non-random distribution of their independent breakpoints, which are clustered in what we called “potentially fragile regions” (PFRs). Especially prominent was PFR1, where seven out of the 30 segmental aneuploidy breakpoints we detected fell within a 3.7 Mb region. Interestingly, this region coincides with a hotspot for genome content variability within *A. thaliana* ^65^ and PFRs overlap three of five regions reported as being particularly prone to rearrangements across *A. thaliana* accessions^68^. Additionally, a Chromosome 1-derived ring mini-chromosome has been reported with a breakpoint at the core of PFR1, with further rearrangements heavily reusing this site ^69^. These observations suggest that its fragility is relevant also in diploids and beyond just the L*er* accession.

PFR1 and PFR5 overlap with two of the three ancient end-to-end fusions that reduced the chromosome number from eight to five in the *A. thaliana* lineage, suggesting that over the millions of years since their origin, these sites have remained fragile within *A. thaliana*. This is similar to what has been reported in the grass, *Phleum echinatum*, where intra-chromosomal telomere repeat regions, presumably footprints of ancestral fusions that also differ among species, show evidence of persistent fragility ^37^. In fact, PFR1 and PFR5 correspond to some of the regions involved in most rearrangement breakpoints in the Brassicaceae more broadly (based on rearrangements reported in ^43^). This correlation connects with the questions: what makes these sites fragile and why do they remain so, despite their costs? Unfortunately, these questions are not fully answered, but there are intriguing hints. It has been long recognised from work in animals that the reuse of fragile sites across species means their fragility is conserved, hinting that their fragility is either directly functional important, or that they are structurally constrained such that selection cannot work effectively against their fragility ^29,52^. In animals, fragile sites tend to replicate late, can lie at chromatin domain boundaries, are particularly prone to replication stress, and can have topological tension ^29,52,70–72^. This would be consistent with our observation that PFR5 is AT enriched, as these sequences can be linked to replication fork stalling ^30^. Replication timing has been mapped in *A. thaliana* ^73^, but at least at the broad scale, there is no link between the PFRs and regions of late replication timing, though PFRs nevertheless may inhibit replication forks in some other way. Another potentially interesting link is that fragile sites in animals are often nuclear lamina associated ^74^. Interestingly, a site recurring multiple times as a deletion breakpoint in mutagenesis in *A. thaliana* also shows evidence of nuclear lamina association ^35^. Our evidence suggests that different fragile sites may be fragile for different reasons, but the idea that their fragility may link to important nuclear functions and 3D genome organization could help explain why they are maintained despite their costs.

PFR1 and PFR5 also overlap with large clusters of nucleotide-binding and leucine-rich repeat (NLR) genes, which are important for pathogen responses ^61,75–77^. NLR genes are frequently in large tandem repeat clusters that have been associated with intra-specific structural variation, likely due to unequal recombination and other meiotic errors associated with repetitive sequences ^8,59–61,78,79^. Thus, it is intriguing that two fragile sites overlapping hot regions for structural rearrangements within the *Brassicaceae* are associated with NLR clusters. Could association with fragile regions pave the way for the frequent rearrangements in these clusters, e.g. via changes in the number and complexity of immunity-related genes ^8,59,60^? Or does the complexity of the NLR clusters directly cause fragility? The causal relationship remains to be determined, but since not all NLR clusters seem to be fragile, we favour the first possibility, that NLR gene cluster diversification is a consequence, not a cause, of fragility in these regions (though we cannot rule out that the association is coincidental).

Collectively, our work supports the hypothesis that the association between chromosome fragility and chromosomal rearrangements is not only a feature of animal systems, but is conserved in plants as well. Overall, this suggests that *de novo* breakage may be a strong driver for genome evolution. This becomes particularly potent in the context of polyploidy, where aneuploidies are less strongly selected against. Preferential breakage at fragile sites may contribute a degree of non-randomness to the deletions and re-arrangements that arise after polyploidy.

## STAR METHODS

### 1. EXPERIMENTAL MODEL AND SUBJECT DETAILS

We produced *Arabidopsis thaliana* neo-polyploids using mutants that yield unreduced gametes (*tam-3*, tam*-4, osd1-3, ps1*) ^80–82^. This allowed us to analyze the progeny of neo-tetraploid inbred individuals right after polyploidization while avoiding ploidy chimeras that often resulting from colchicine treatment. We crossed *tam-4* (CSHL_ET12273) and *osd1-3* (CSHL_ET1227) homozygous mutants (in L*er*), which yield unreduced gametes, to generate double heterozygous neo-tetraploids ^82^ with restored reductional meiosis (Figure S1). Similarly, we used *tam-3* (SALK_080686) and *ps1* (SALK_078818) mutations in Col-0 ^80,82^ to generate neo-tetraploid Col-0 inbreds (Figure S1). We generated bi-directional crosses (three in each direction) between the neo-tetraploid Col-0 and neo-tetraploid L*e*r individuals to generate F_1_ hybrids (Figure S1A). The genome content and euploidy of the neo-tetraploid Col and L*er* parents was verified using flow cytometry on DAPI-stained leaf nuclei, as well as by sequencing of their leaf DNA (see below). A total of 228 F_1_ hybrids were sown on soil for three weeks, and the leaves of each individual were collected for DNA extractions. The hybrid genotype of the F_1_ individuals was verified by checking that they are heterozygous for published SSLP markers ^83^.

### 2. METHOD DETAILS

#### DNA extraction

Genomic DNA was extracted using a protocol modified from ^84^. Briefly, 0.1-0.3g of liquid nitrogen frozen leaf tissue was homogenized using a TissueLyser II. Tissue was then lysed in a lysis buffer (100mM Tris-HCl pH8, 500mM NaCl and 1.3% SDS) supplemented with RNase A and incubated at 55°C for 30 min. After incubation, the supernatant was collected by centrifugation and mixed with 0.325 volumes of 5M Potassium acetate on ice for 5 minutes. DNA in the cleared supernatant was bound to 1 volume of homemade Sera-Mag magnetic bead solution (0.4% washed hydrophobic carboxylate-modified beads, 11% PEG 8000, 1.6M NaCl, 10mM Tris-HCl pH8 and 1mM EDTA) for 10 minutes. The beads were magnetized, washed twice with 80% ethanol, air-dried, and the DNA was eluted in 50 μL of pre-warmed nuclease-free water.

#### Library-preparation and sequencing

For the parental neo-tetraploid lines, library-preparation was performed using a custom protocol. In brief, 50 ng of genomic DNA in a 9μl volume was digested using 1μl of Illumina Tagment TDE1 enzyme, and 10 μl of Illumina 2X Tagment TD buffer, and incubated at 37 <u>°</u>C for 30 minutes on a heat block. The resulting fragmented DNA was immediately cleaned up using column-purification with the Zymo DNA Clean & Concentrator kit and eluted in 11μl of 10mM Tris HCl. 3 μL of the eluate was used as a template for a PCR reaction (12 cycles) incorporating Illumina indices, followed by a 1.8:1 ratio cleanup using AMPure XP magnetic beads. The 6 resulting genomic DNA libraries were subsequently sequenced at an average read coverage of 15X at the Functional Genomics Centre Zurich (FGCZ).

For F1 hybrids and control samples, library preparation and whole genome-sequencing (paired-end, 150 bp sequencing reads) was performed by Novogene UK in two batches, at an average coverage of 22X across all samples (with respect to the TAIR10.1 reference genome). The first batch contained 47 samples (including some diploid controls; 1 Col-0 WT, 1 Ler WT and 1 euploid hybrid) and the second batch contained 190 samples (including diploid controls; 2 Col-0 WT, 2 Ler WT and 2 euploid hybrids).

Read counts for all samples are presented in Supplemental Data S4.

#### Read-mapping and aneuploidy detection through coverage-based approach

This pipeline was a modified version of the pipeline used in ^7^. We first created an *in silico* hybrid reference genome using five chromosomes of TAIR10.1 and five chromosomes of Ler ^66^. Read1 and Read2 were aligned independently to the hybrid reference genome using *bwa-mem* (in single-end mode, default options) and filtered to retain SAM flags 0 and 16. The sam files were subsequently merged and converted to .bam files using samtools. Reads carrying XA and SA tags were then removed from the .bam files to generate a .bam file having uniquely mapped reads. Reads mapped to each of the TAIR10.1 and Ler subgenomes were then separately analysed for the number of mismatches they exhibited (from 0-5). Based on the observed mismatch numbers in all of our samples, we retained reads with 0-2 mismatches in the TAIR10.1 genome and reads with no mismatches in the Ler genome. All of these filtering steps ensured we had largely subgenome-specific reads, but this stringency resulted in the retention of only ∼10% of the initial sequencing reads. Mapping statistics for all samples are presented in Supplemental Data S5.

The filtered .bam alignment files were then sorted with *samtools sort*, followed by removal of PCR duplicates using *picard*. The number of aligned reads at 100kb windows throughout the *in silico* hybrid genome (non-overlapping) were then computed using *bedtools* coverage (bedtools version 2/2.31.0). The read counts for all samples (across all windows) were combined into a single file using bedtools unionbedg. This composite file was then modified in R, as follows:

1. Windows were first split into TAIR10.1 and Ler subgenomes respectively
2. For every sample, outlier windows were identified as those that exhibited read counts more than that of 98^th^ percentile of all windows, or less than 2^nd^ percentile of all windows. These outlier windows were changed to “NA”
3. Read counts per window were then normalised to the total sum of read counts across all windows (for a given sample) and excluding the “NA” values. This normalised read count will be referred to as “value [1]”.
4. For each window, the mean of value [1] across three euploid hybrid controls was calculated. The three controls were samples “Col2BXLer3D”, "HyB_A " and “HyB_B”. This mean value will be referred to as “value [C]”
5. For each window, normalised read coverage was calculated as (value [1] / value [C])* 2 (since each subgenome should have 2 copies in a euploid).

Normalised read coverage for each window was then plotted across all chromosomes from both subgenomes, for each sample, using R. Samples with aneuploid chromosomes or aneuploid chromosomal segments were identified first by visually examining the plots for significant deviations of normalised read coverage from 2, with increases denoting “gains” and decreases denoting “losses” of chromosome/segmental copies.

We also independently verified the aneuplodies also using a custom python-based pipeline described in ^38^, where we used our filtered reads (.sam alignment files) as an input for the pipeline, and chose one of the euploid Hybrid samples (“HyB_A”) as a control.

#### Read-mapping and aneuploidy detection through variant-calling approach

Since the coverage-based approach (above) utilized only a small fraction of all sequencing reads, we complemented it with a variant-calling approach to detect aneuploidy, similar to the methods used in ^39^. In this approach, Read1 and Read2 of every sample were mapped in single-end mode to the both the (1) Ler reference genome and (2) TAIR10.1 reference genome using *bwa mem* (default options) and filtered to retain SAM flags 0 and 16.

The .sam files were subsequently merged and converted to .bam files using samtools. Reads carrying XA and SA tags were then removed from the .bam files to generate a .bam file having uniquely mapped reads. This .bam file was sorted using samtools, and *picard* was used to remove read duplicates and add new read groups. The resulting .bam file was then subjected to a standard variant calling workflow using GATK (v3.8), where only high quality bi-allelic variants were retained. Mean allele frequencies of the non-reference variants at 10kb and 100kb windows (non-overlapping) across the genome were calculated using *bedtools* map. The mean allele frequencies were plotted in R for visualisation and subsequently used to determine segmental aneuploidy breakpoints (see below). Mapping statistics are presented in Supplemental Data S6 (mapping to TAIR10.1 reference) and Supplemental Data S7-S8 (mapping to Ler reference).

#### Mapping of segmental aneuploidy breakpoints and identification of PFRs

From both the coverage-based method, and the variant calling method with 100kb windows, we initially obtained rough estimates of the positions corresponding to segmental aneuploidy breakpoints. To have a better resolution of identifying the breakpoints, we calculated the mean non-reference allele frequency in 10 kb windows and manually inspected the candidate regions for stark drops/gains in non-reference allele frequencies where a segmental loss/gain was expected. This helped to identify 10 kb intervals where the aneuploidy was first observed and the last interval in which the non-reference allele frequency remained in the same range, before exhibiting a drop/gain. The upstream coordinate of the first interval and the downstream coordinate of the last interval were then chosen as the “start” and “end” positions of the segmental aneuploidies.

For each of the above start/end positions that was not close to the start or the end of a given chromosome, we calculated a window of 250kb upstream and 250kb downstream. Each such window was assigned as the coordinates of a breakpoint. In this manner, we found a total of 21 segmental aneuploidies and 30 segmental breakpoints.

From both the coverage-based method, and the variant calling method with 100 kb windows, we initially obtained rough estimates of the positions corresponding to segmental aneuploidy breakpoints. To improve the breakpoint resolution, we calculated the mean non-reference allele frequency in 10kb windows and manually inspected the candidate regions for stark drops/gains in non-reference allele frequencies where a segmental loss/gain was expected. This helped to identify 10 kb intervals where the aneuploidy was first observed and the last interval in which the non-reference allele frequency remained in the same range, before exhibiting a drop/gain. The upstream coordinate of the first interval and the downstream coordinate of the last interval were then chosen as the “start” and “end” positions of the segmental aneuploidies.

In all cases where the aneuploidy start/end positions did not overlap with the start or the end of a given chromosome, we calculated a window of 250kb upstream and 250kb downstream. Each such window was assigned as the coordinates of a “**breakpoint”**. In this manner, we identified 30 segmental breakpoints when mapping reads to the TAIR (Figure S9) genome and 30 segmental breakpoints when mapping reads to the Ler genome. From these, we further identified five potentially fragile regions (PFRs) where breakpoints were recurrent.

For both TAIR10.1 and L*er* reference genomes, the coordinates of the breakpoints and PFRs differed (likely due to genome size differences, with the TAIR10.1 genome being 119 Mb and Ler genome being 115 Mb). However, the overall PFR positions relative to centromeres in each chromosome were largely similar.

All sequencing data mapping, variant calling and aneuploidy detection pipelines were carried out using the EULER high-performance compute cluster of ETH Zürich.

#### Precise mapping of ancient fusion in *A. thaliana* Chr1

To precisely map the breakpoint of the ancient fusion that gave rise to *A. thaliana* Chr1 we selected a set of anchor genes in the studied region in TAIR10.1 reference and registered the orthologs in six related species: *Arabidopsis haleri*, *Arabidopsis arenosa*, *Arabidopsis lyrata*, *Capsella rubella*. We chose anchor genes with single copy unambiguous orthologs in the mentioned species according to Jbrowse 2 (www.arabidopsis.org, Figure S6) or our own annotations (in case of *A. arenosa*). We included an additional species (*Cardamine hirsuta*), using synteny blocks (identified in ^85^). Next, we retrieved the chromosome identities for all the anchor genes and synteny blocks map (Figure S6) for each species and compared them. With this, we compared for each anchor gene on which chromosome they map to for each ortholog in all the species to infer the breakpoint. Since we observed a complex array of rearrangements (fusion, translocation and inversion) whose chronology could not be inferred. Nevertheless, we assigned the precise fusion breakpoint somewhere at the end of the northernmost anchor gene of the ancestral Chr1 that neighbours an *A. thaliana* gene located in the ancestral Chr2. Finally, we used the coordinates of the L*er* orthologs to map the breakpoint in L*er* (shown in Figure 3B) for positional comparison with the *de novo* breakpoints.

#### Analysis of evolutionary breakpoints in Brassicaceae family

We used the data from ^43^ that identified ancient rearrangements between 65 conserved syntenic blocks in ten species within the Brassicaceae family: *Arabidopsis lyrata*, *Arabidopsis thaliana*, *Erysimum cheiranthoides*, *Megadenia pygmaea*, *Thlaspi arvense*, *Arabis alpina*, *Draba nivalis*, *Meniocus linifolius*, *Tetracme quadricornis*, and *Aethionema arabicum*. We used the order of synteny blocks for each chromosome as well as for the corresponding tree nodes for each of the branches of the phylogenetic tree available in ^43^. We made a correction in the original data: we detected that some blocks that were previously detected as inverted in the original synteny blocks (D, P R, Q and S ^41^), the updated version ^43^ contained subdivisions of those blocks with coordinates in inverse order (e.g. R5, R4, R3, R2, R1, instead of R1, R2, R3, R4, R5). The corrected list of 65 blocks is available in Supplemental Table S14. We verified this corrected order using the data from an independent synteny study ^85^.

We counted, for each of the 65 conserved synteny blocks, how many different neighbour blocks they had (in their north and south boundaries) across the different species and nodes. For each block boundary (north and south) we subtracted one to the count of neighbours to obtain the final number of times that each block boundary participated in a rearrangement. The original data used TAIR gene annotations as references. We transferred these annotations to L*er* reference using the ortholog gene (annotations in ^66^. When a TAIR gene was not present in the L*er* assembly the closest gene was used instead. Then, we retrieved the coordinates of the start and stop of the first and last *A. thaliana* genes (L*er* reference,), respectively, of each block and used them as evolutionary breakpoints when they have been involved in any rearrangement. Finally, we plotted for each 1 Mb bin the number of uses as an evolutionary breakpoint of the overlapping block boundaries and calculated the genome-wide average and the first and fourth quartiles as threshold criteria to identify the hot and cold regions, respectively, for ancient rearrangements (Figure 3B).

#### Custom TE annotation of the Ler genome

The Ler genome reference and annotation ^66^ was downloaded from NCBI (GenBank assembly accession: GCA_001651475.1). To identify repeat regions and Transposable elements (TEs), we used EDTA (https://github.com/oushujun/EDTA; ^86^. For LTRs that were annotated based on structural homology, we retained only the parent feature. TE sequence ontologies provided in the EDTA output files were used to infer the TE superfamilies of each element.

### 3. QUANTIFICATION AND STATISTICAL ANALYSIS

#### Criteria for aneuploidy assignment

We only scored aneuploidies when we confirmed that shifts in coverage were concomitant with complementary changes in allele frequency. When two different chromosomes were affected in the same aneuploid individual, we counted them as different aneuploidies. Conversely, when two or more different fragments were affected by segmental gains or losses in the same chromosome from the same parent we counted them as part of the same segmental aneuploidy.

#### GO and NLR enrichment analysis

We selected all the genes present in all the hot and cold regions using the annotations in Ler genome ^66^. Next we performed GO enrichement analysis using the PlantTregMap online tool annot https://plantregmap.gao-lab.org ^87,88^.

To analyze the enrichment of NLR genes within PFRs we switched to the higher quality TAIR assembly where the highly repetitive NLR paralogs are more accurately annotated, we used the PFR coordinates based on mapping breakpoints to TAIR genome given the consistency with the mapping on L*er* (Figure S8).

#### Genomic and epigenomic properties of breakpoints and PFRs

To test whether the genomic/epigenomic properties of breakpoints and PFRs were significantly different from other genomic regions, we separately examined two sets of data (1) all breakpoints (2) each of the five PFRs individually.

For each table, we first generated a set of 5000 random regions corresponding to the same size intervals as the “test” table (eg. breakpoints/PFRs) and excluding them from the sampling. This was carried out using *bedtools shuffle.* For both “test” and “random” regions, we then examined:

1. Number of overlapping genes using *bedtools closest*
2. AT/GC content (using *bedtools nuc*)
3. Number of overlapping TEs (total), and number of overlapping TEs in each superfamily (where superfamily refers to TE sequence ontologies from EDTA output data), using *bedtools closest*
4. Chromatin accessibility properties of Ler: using Ler-1 wildtype accessibility levels in counts per million (mean of 3 biological replicates) from 31,295 accessible chromatin regions identified from 18 *A. thaliana* accessions and their *met1* methylation mutants; Srikant et al. 2022 (mean Ler-1 accessibility level was examined across all dACRs that overlapped a given PFR/breakpoints).

For (4), we could only use the breakpoints and PFRs derived from TAIR10.1 mapping since the original table from ^89^ was mapped to the TAIR10 reference genome. All other features were examined using both TAIR10.1 and Ler reference genomes.

To identify whether a given feature was significantly different between the “observed region” (eg. PFR1-5, Breakpoints) and the mean of all randomly sampled regions, we carried out the following steps:

1. We first used a two-tailed test to identify the p-value, where p.value=mean(abs(permuted_values) >= abs(observed_value)). In cases where the p-value was >0.95 (alpha=0.05), we applied a one-sided test where p.value=mean(abs(permuted_values) <= abs(observed_value)). This additional test helped us identify whether certain features were lower in magnitude than expected by chance.
2. We then applied a multiple-testing correction for all the features tested (Benjamini Hochberg) to obtain adjusted p-values. This was carried out separately for each of the PFRs, and also for the set of breakpoint regions.

#### Statistical analyses

All statistical tests and p-value correction (except for genomic feature analysis) were performed using R studio. The results from all tests, including hypotheses tested, tests used, p-values, and p-value correction methods are available in Supplemental Table S18 for genomic feature analysis and S19 for the rest.

### 5. KEY RESOURCES TABLE

#### Key resources table

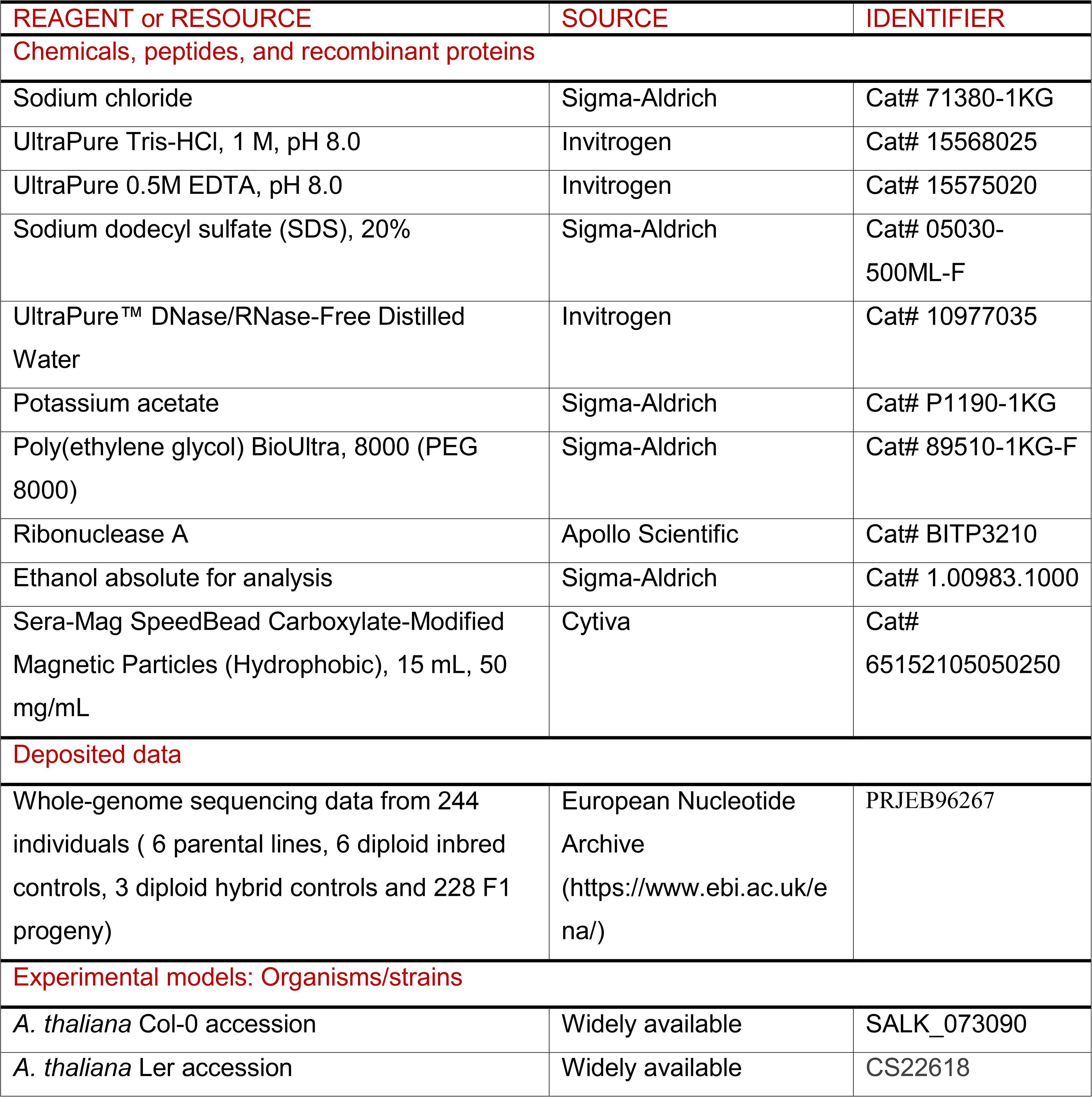
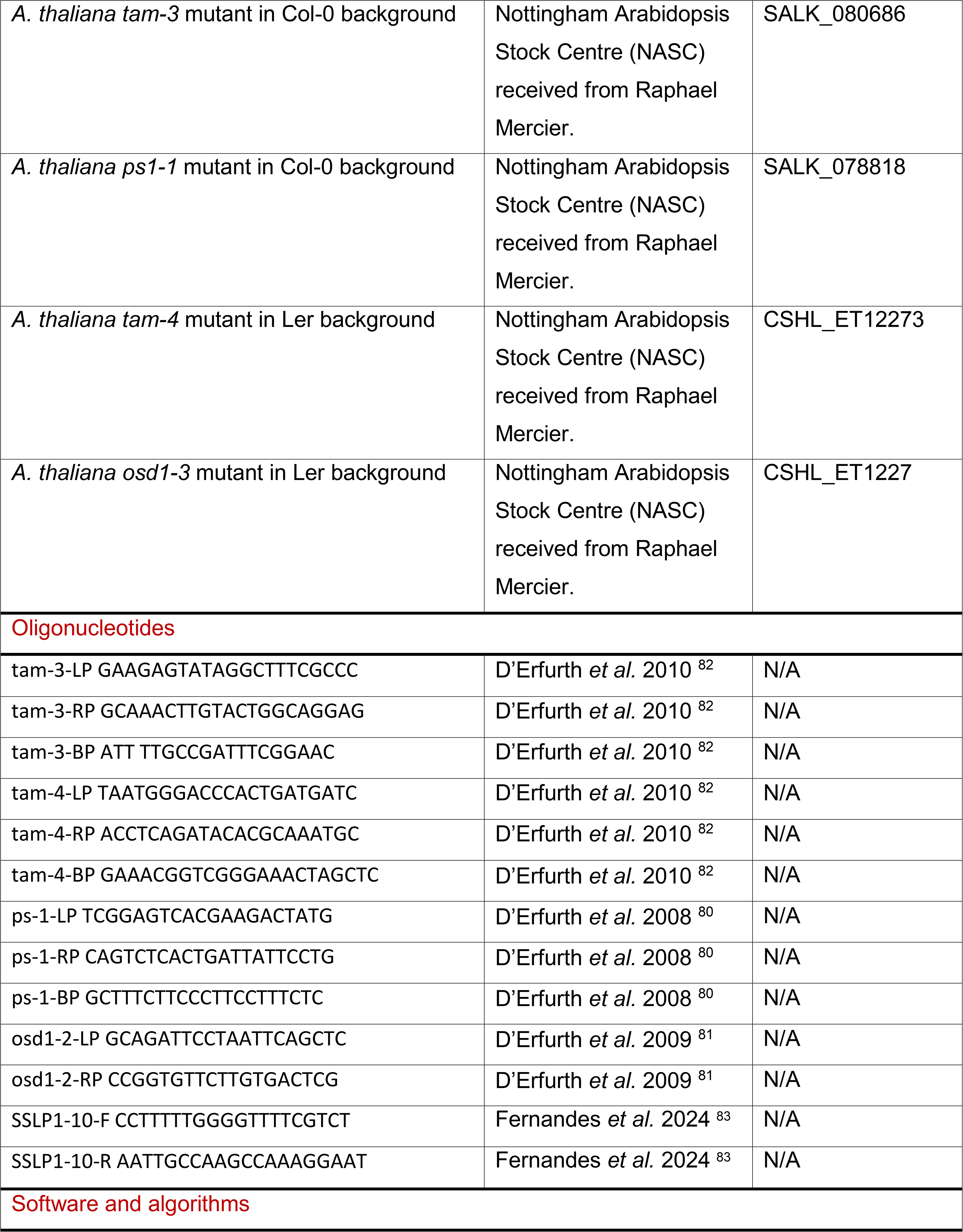
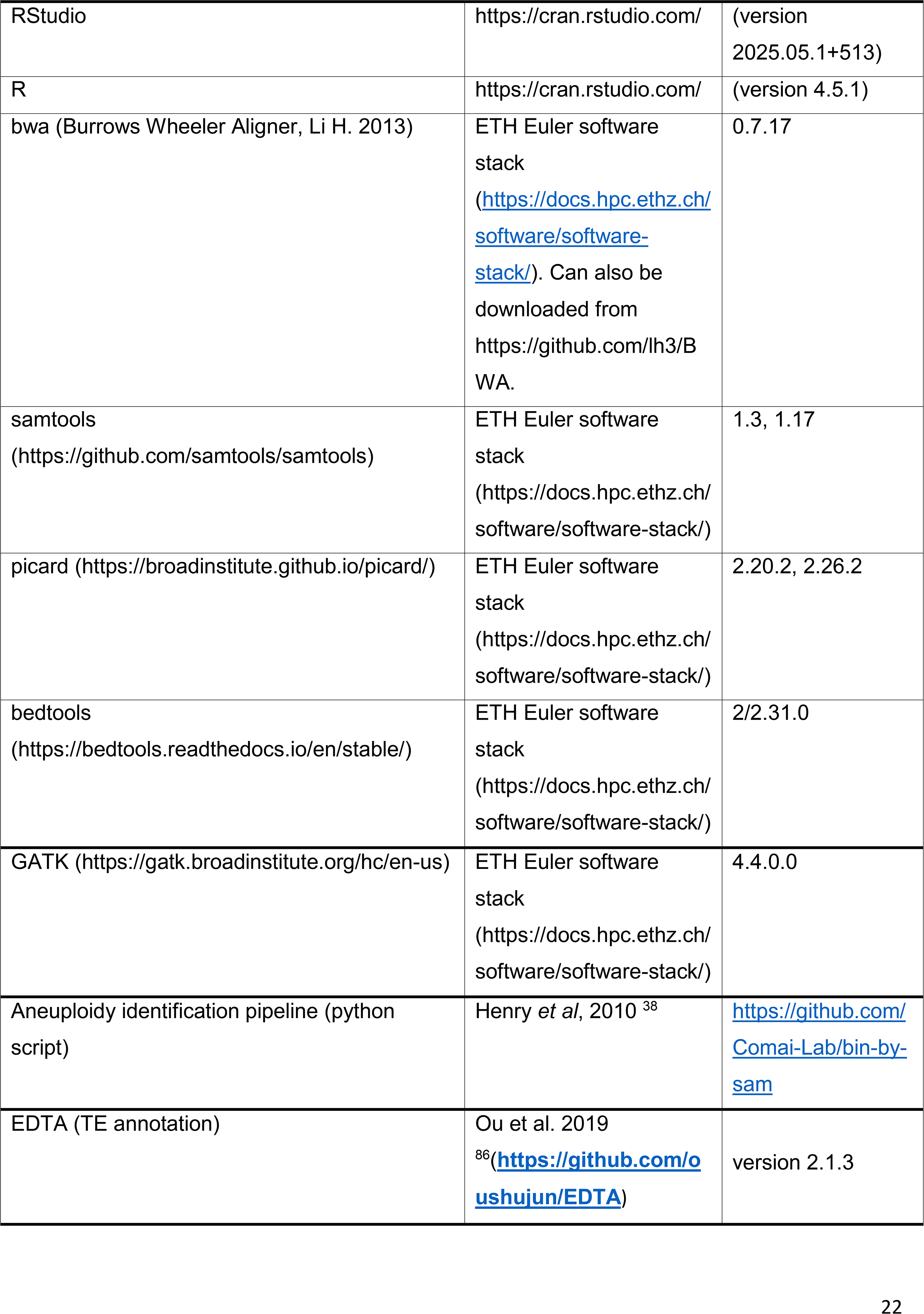
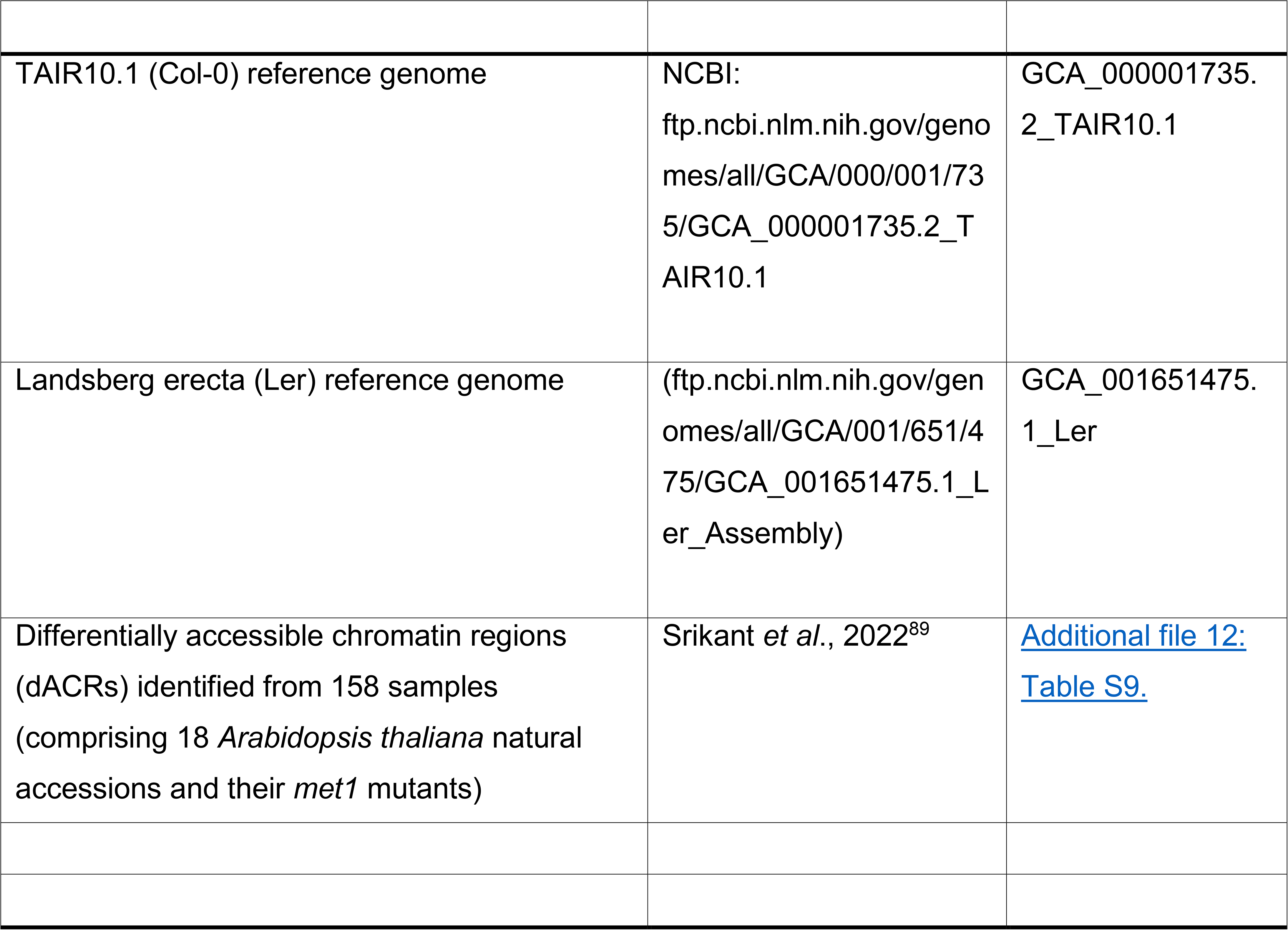

## ACKNOWLEDGEMENTS

This project was funded by the project ProSPECT within the MSCA-IF-2020 program (European Commission, grant number 101029732), a Career Seed Award from ETH Zürich to A.G. and a Swiss National Science Foundation Ambizione Grant awarded to T.S. (Grant number: 233372) and core funds from ETH-Zürich to K.B. We thank Nora Walden and Jie Liu for facilitating access and clarifications about their synteny data, Raphael Mercier for providing seeds for mutants with unreduced gametes, Oriol Garcia Prats for support producing plant materials, and Rajeev Kumar for scientific discussions. We thank the Functional Genomics Centre of Zurich, and the ETH’s Genetic Diversity Centre for providing access and training to use their facilities.

## AUTHOR CONTRIBUTIONS

AG conceptualized the research; AG and HST generated the plant materials; TS, AG performed the research; TS and AG analysed the data; KB, AG and TS secured funds; AG drafted the manuscript; AG, KB and TS produced the final manuscript.

## Supplementary information

**Figure S1:**
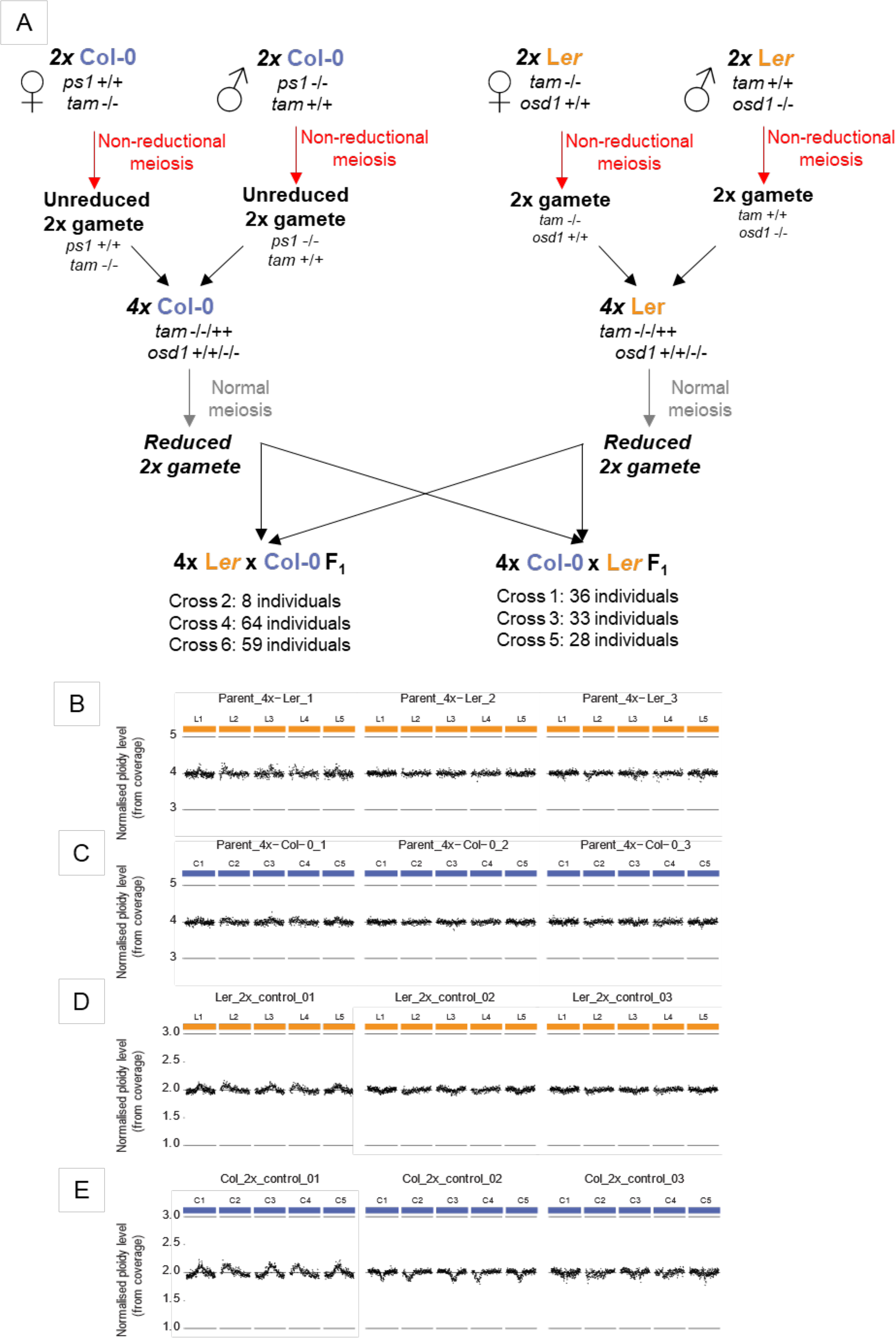
(A) Genetic crossing strategy used to obtain first-generation neo-tetraploids. (B) Coverage plots of three Ler neo-tetraploid inbred parents mapped to the Ler reference genome (orange); (C) three Col-0 neo-tetraploid inbred parents mapped to TAIR10.1 reference genome (blue). (D) three diploid Ler inbred lines mapped to the Ler reference genome and (E) three diploid Col-0 inbred lines mapped to the TAIR10.1 reference genome. Each point represents the mean normalized read coverage for a 100kb window. The overall even coverage for all chromosomes in all samples confirms that the neo-tetraploid parents that were used to generate F1 hybrids are euploid (four copies for each chromosome without detectable segmental aneuploidies) and the diploid inbred controls are euploid as well (two copies of each chromosome).

**Figure S2:**
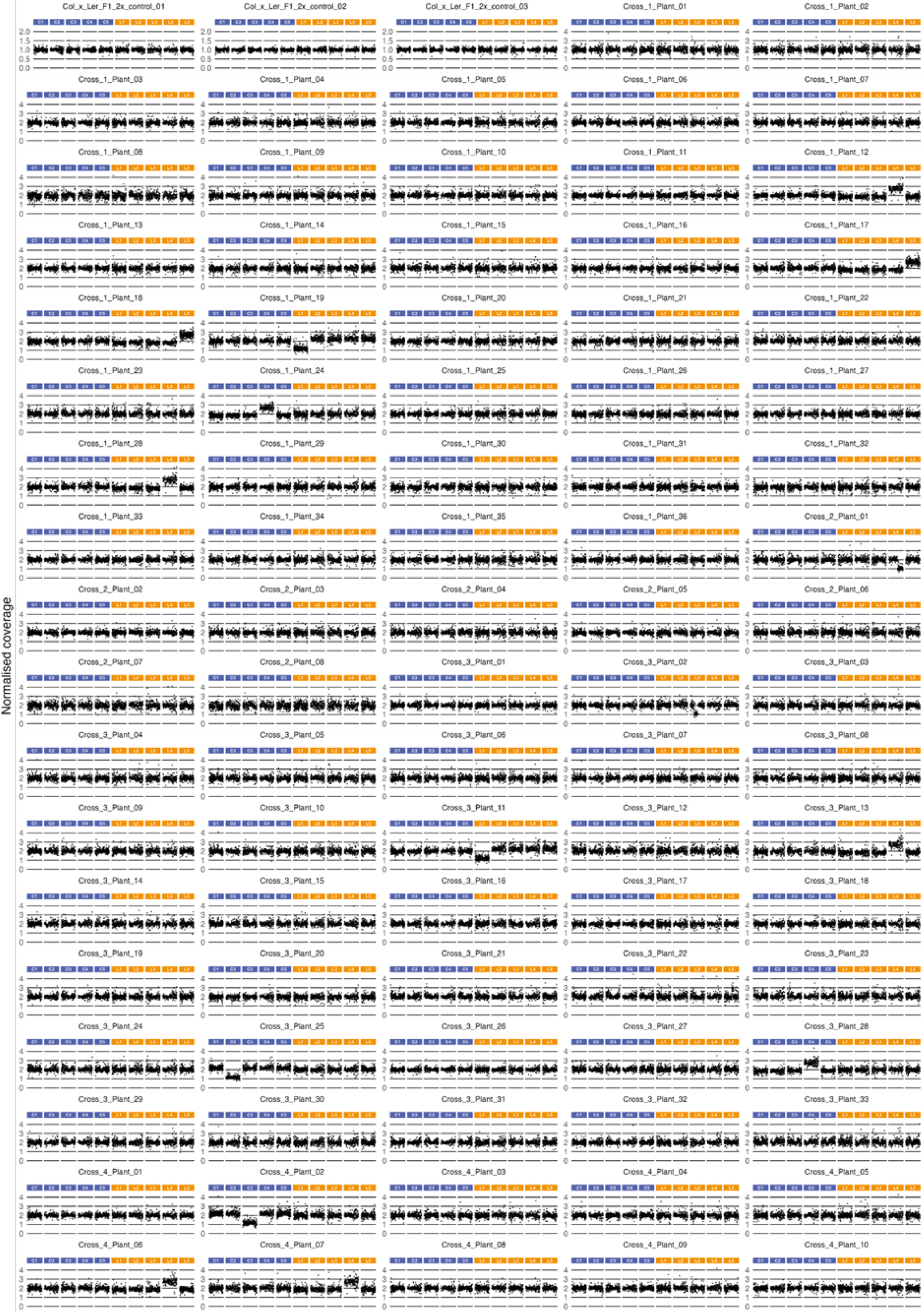
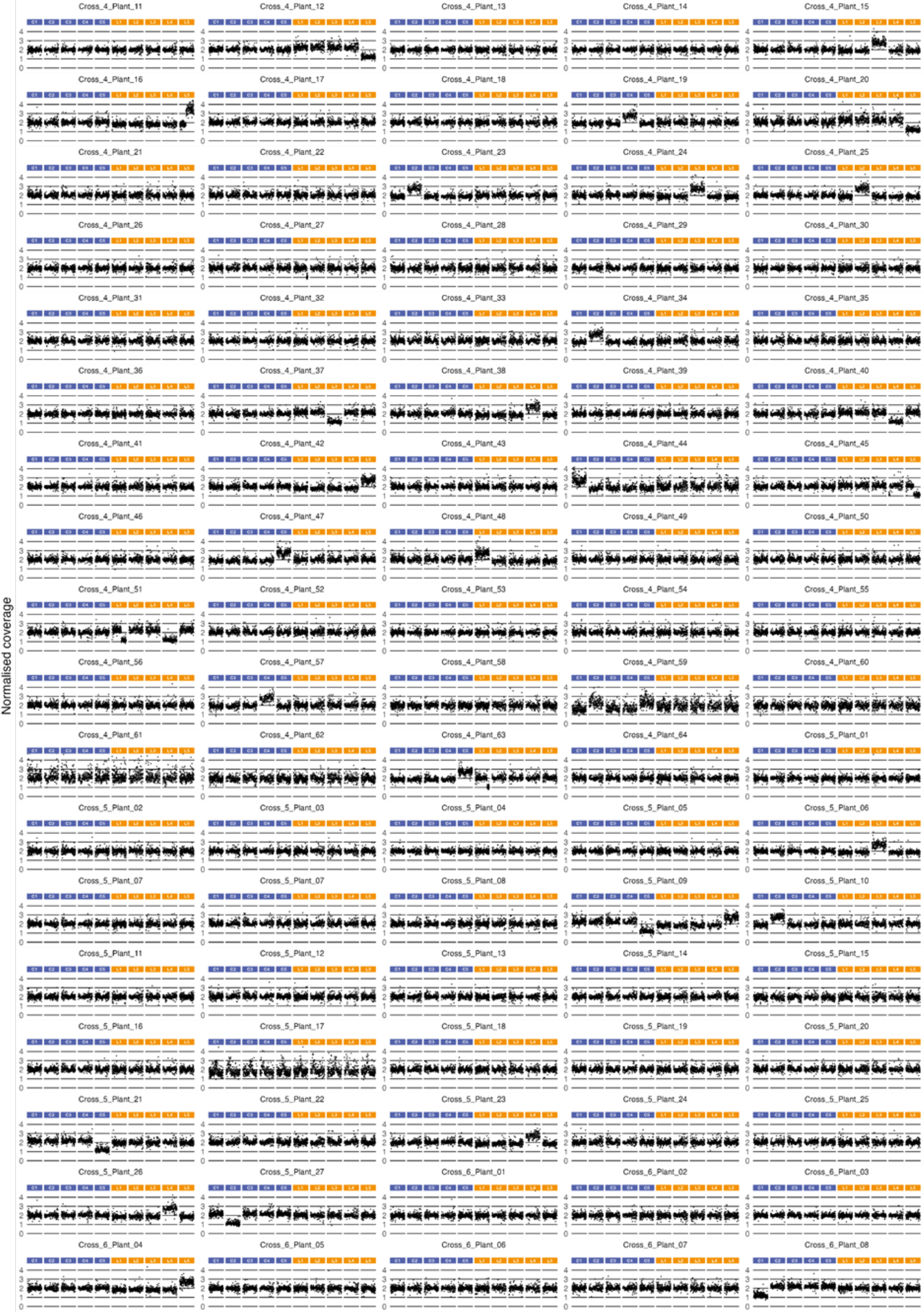
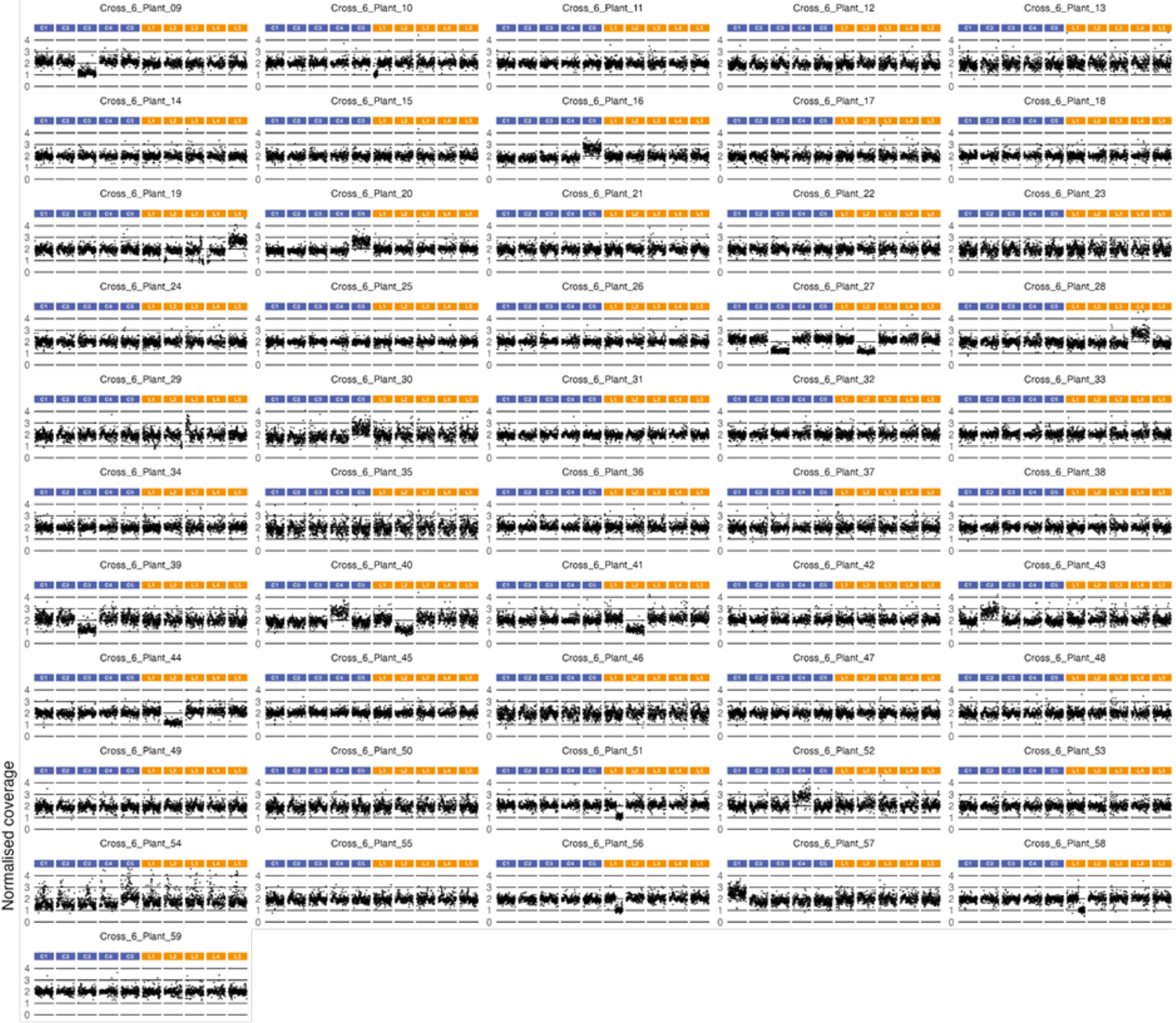
Aneuploidy detection using normalized coverage for all 231 individuals sequenced (228 F1 plants and three diploid F1 hybrid controls). For each sample, a scatterplot of normalized read coverage in 100 kb windows across an *in silico* hybrid genome of Ler and TAIR10.1 is shown. For each subgenome, read coverage at each window was first normalised to total reads and then to the mean read coverage across three euploid 2X hybrid controls of Col x Ler. This value was then multiplied by 2 to generate the relative coverage value, which is plotted in the y axis of every sub-plot. Ler chromosomes are indicated in yellow (labelled L1-L5) and TAIR10.1 (Col-0) chromosomes in blue (labelled C1-C5). Due to the large number of samples analysed, the figure is split into three parts.

**Figure S3:**
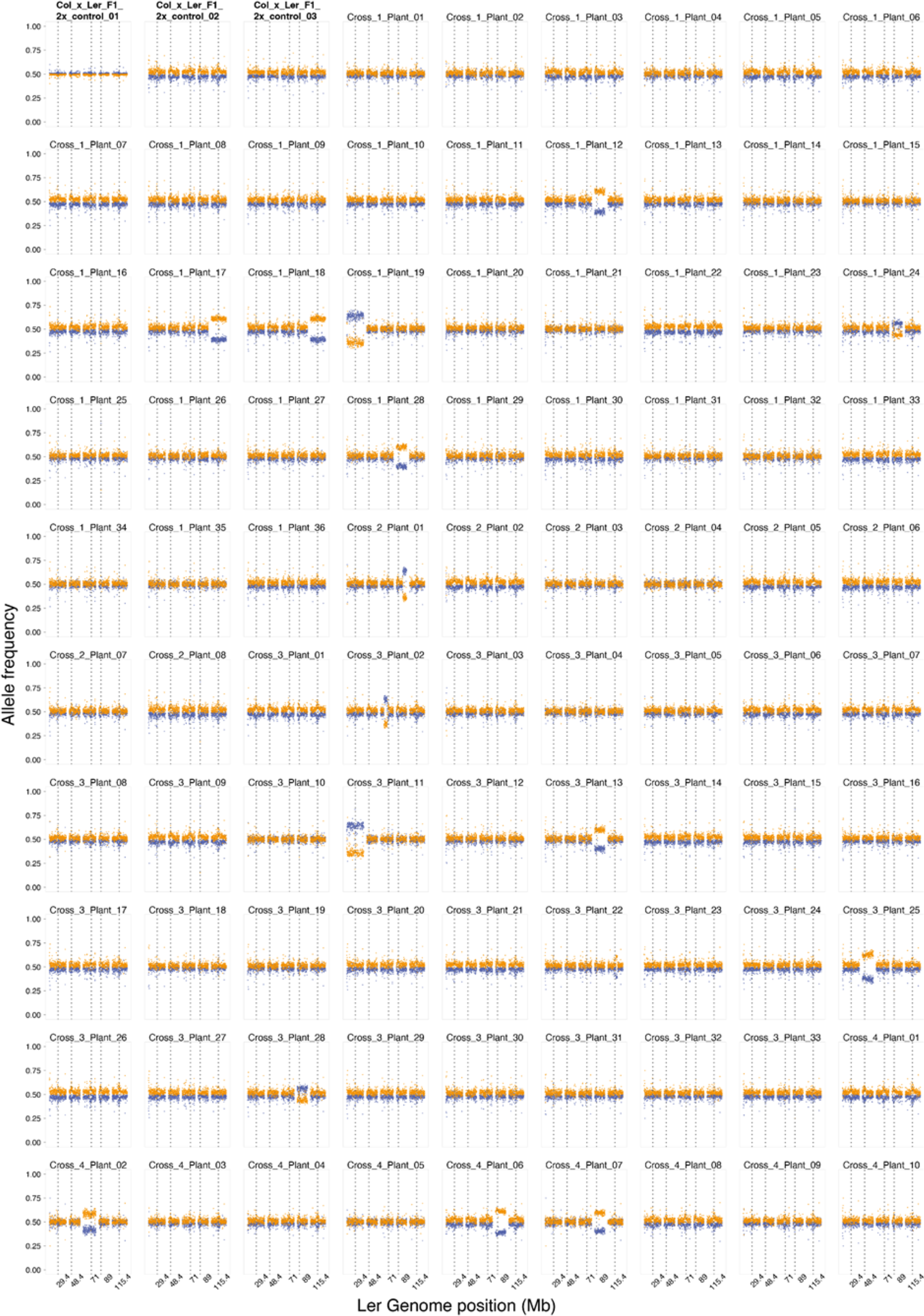
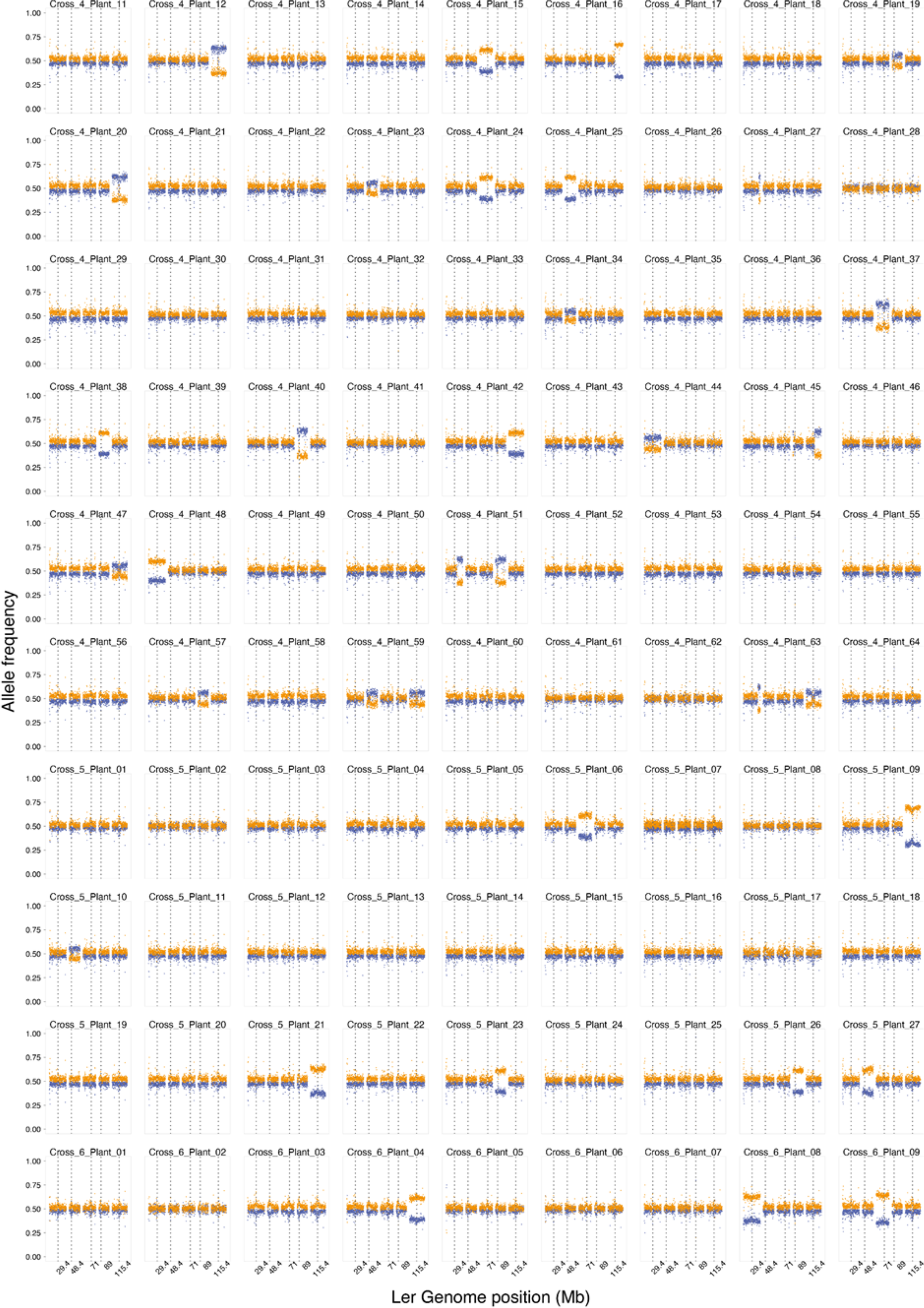
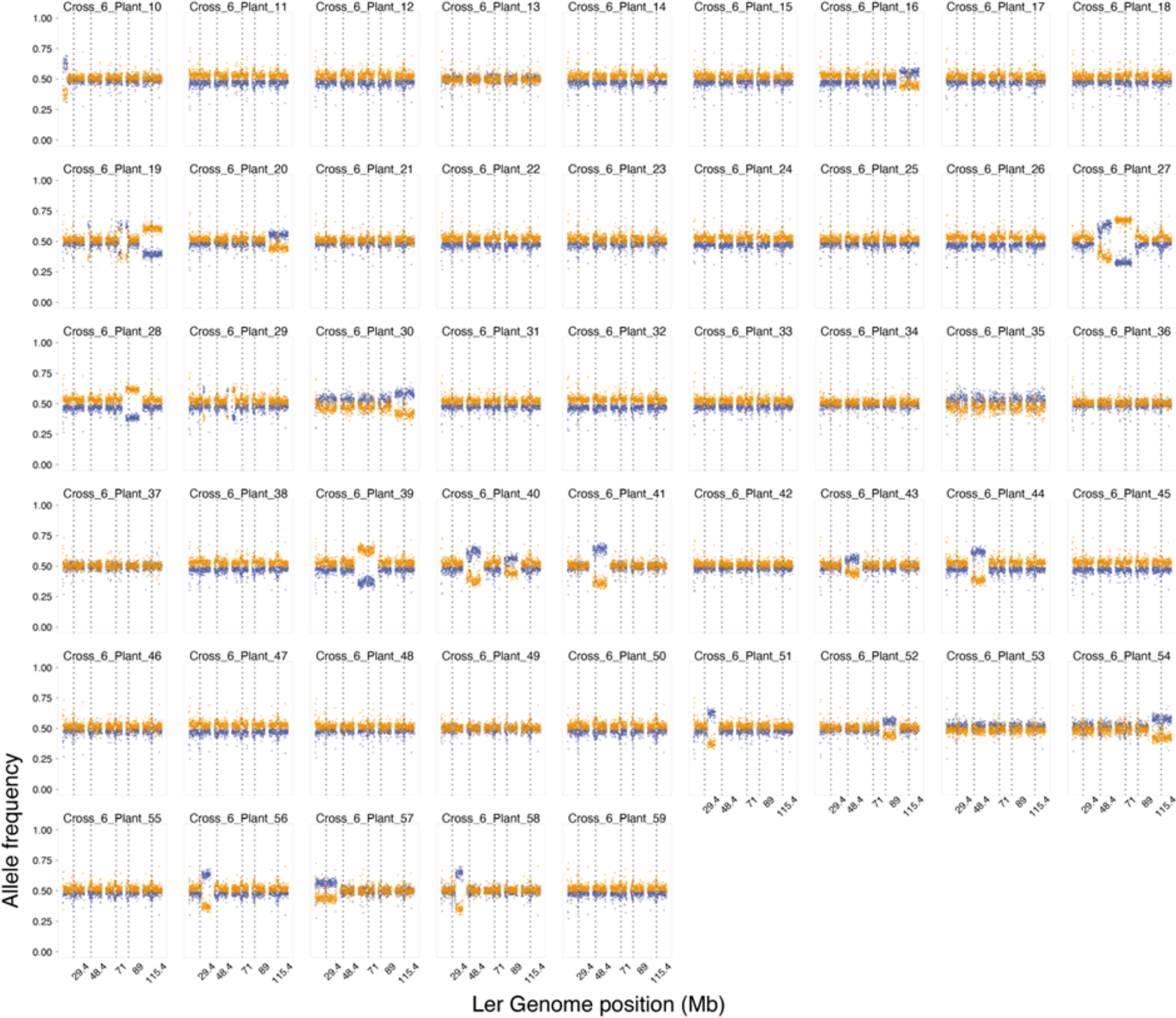
Aneuploidy detection using allele frequencies at bi-allelic variant sites for all 231 individuals sequenced (228 F1 plants, plus three diploid F1 hybrid controls), and mapped to the Ler reference genome. Plots show average reference allele frequencies (Ler, yellow dots) and non-reference allele frequencies (Col-0, blue dots) for 100 kb windows across the genome. Aneuploidies are evident where allele frequencies diverge and form clearly distinct bands at diYerent frequencies.

**Figure S4:**
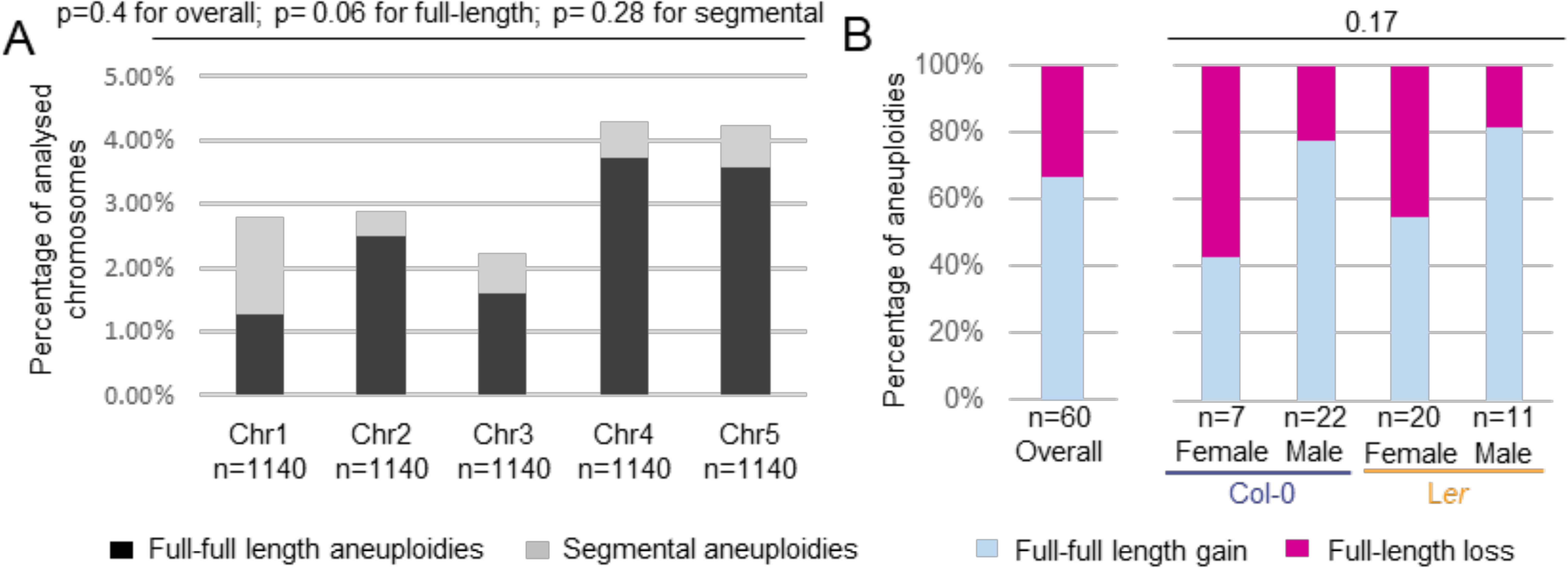
Aneuploidy frequencies for each individual chromosome. (A) Proportion of full-length (dark grey) and segmental (light grey) aneuploidies for each chromosome. The given p-values are from Fisher’s Exact test for differences by chromosome for overall, full-length, and segmental aneuploidies. (B) Proportion of inferred gametes carrying full-length gains and losses for each parent-of origin compared with the overall proportion. The displayed p-value refers to the results from Fisher’s Exact test for chromosome preference for gain/loss proportion, i.e. whether the gain/loss is similar across the samples.

**Figure S5:**
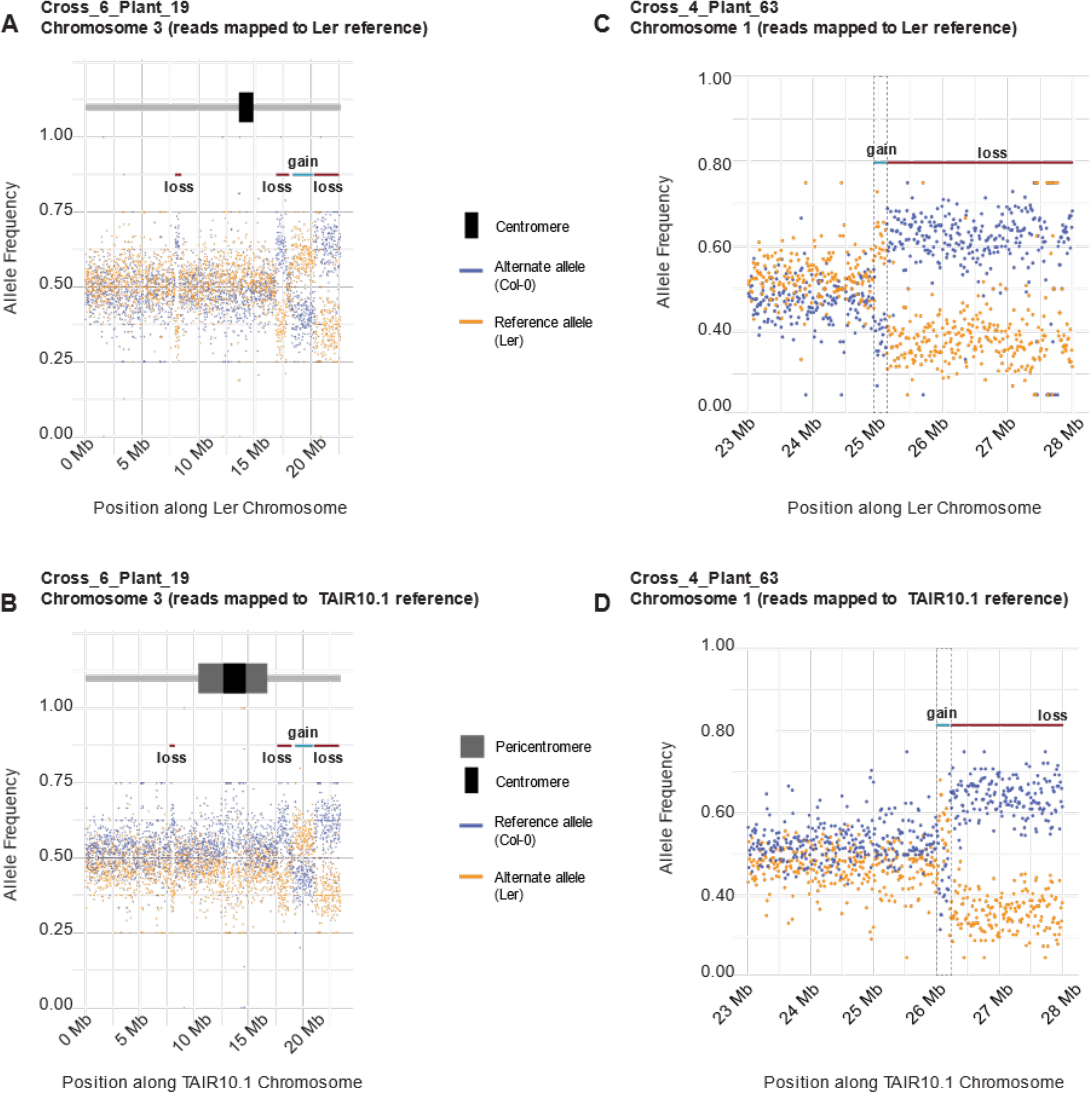
Complex rearrangements identified using the variant-calling approach of aneuploidy detection. (A,B) Chromosome 3 of sample “Cross_6_Plant_19”, which exhibits three segmental losses and 1 segmental gain from Ler. Each dot represents a 10kb window, where mean allele frequencies of Ler and Col-0 alleles were computed when mapping reads to the Ler (A) or TAIR10 (B) reference genome. (C,D) An example of the smallest segmental gain of Ler identified in our dataset, in Chromosome 1 of sample “Cross_4_Plant_64”, when reads are mapped to the Ler reference genome (C) and TAIR10 reference genome (D) respectively.

**Figure S6:**
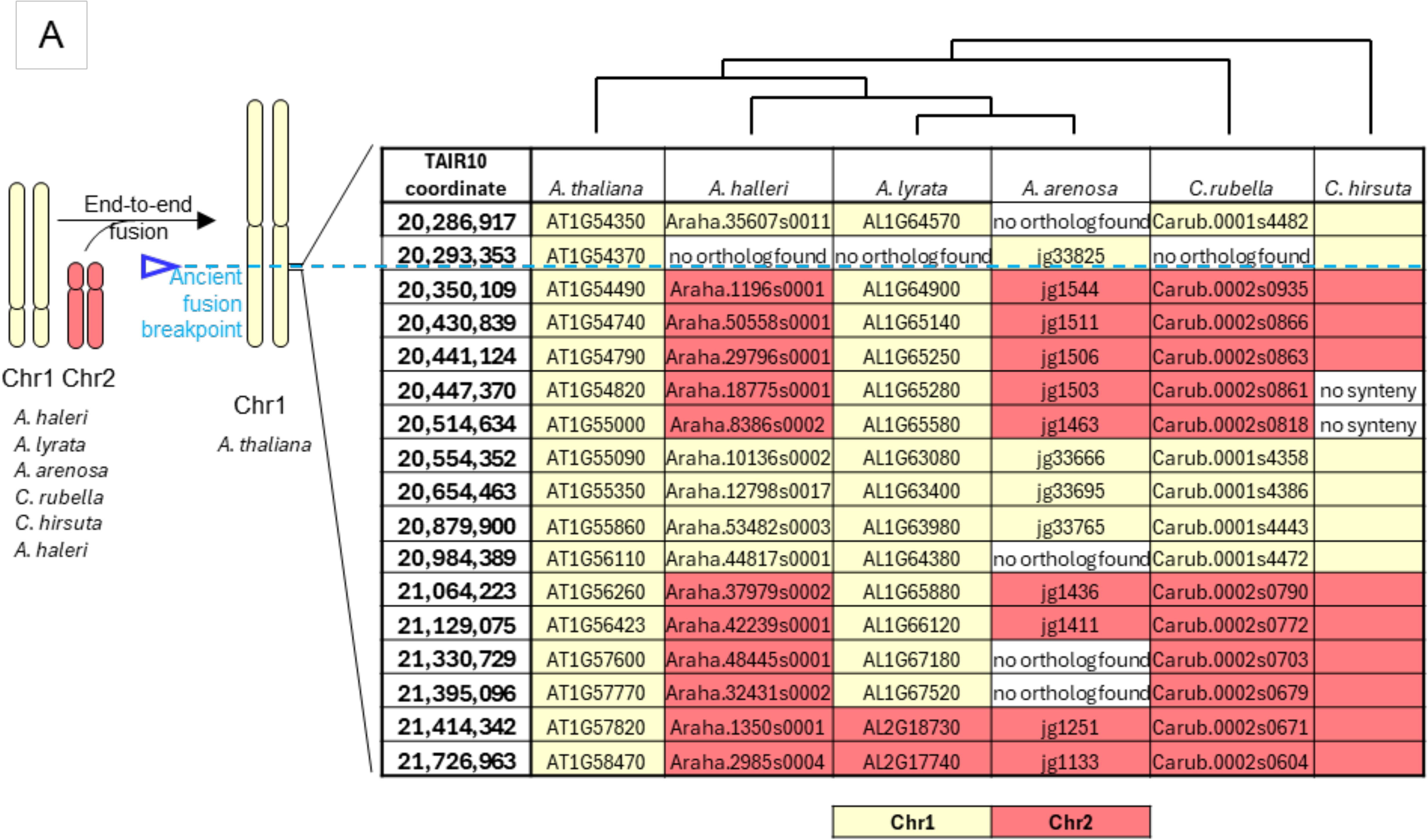
Mapping the breakpoint of the ancient fusion that gave rise to *A. thaliana* Chr1. (A) summarizes the synteny analysis that led to the mapping of the breakpoint. It includes the annotation IDs of multiple anchor genes in the region (in TAIR10 reference; i.e. *Arabidopsis thaliana*), and the syntenic orthologs in the species *Arabidopsis halleri*, *Arabidopsis arenosa*, *Arabidopsis lyrata*, *Capsella rubella* and *Cardamine hirsuta* (see methods). The order of anchor genes reflects their order in TAIR10.1. For each ortholog, the color represents which chromosome it maps to in the respective species, yellow for Chromosome 1 (Chr1) and red for Chromosome 2 (Chr2). The inferred breakpoints in *A. lyrata* slightly diYer from those in the rest of species, potentially due to a previously reported *A. lyrata*-specific translocation or miss-assembly ^1^.

**Figure S7:**
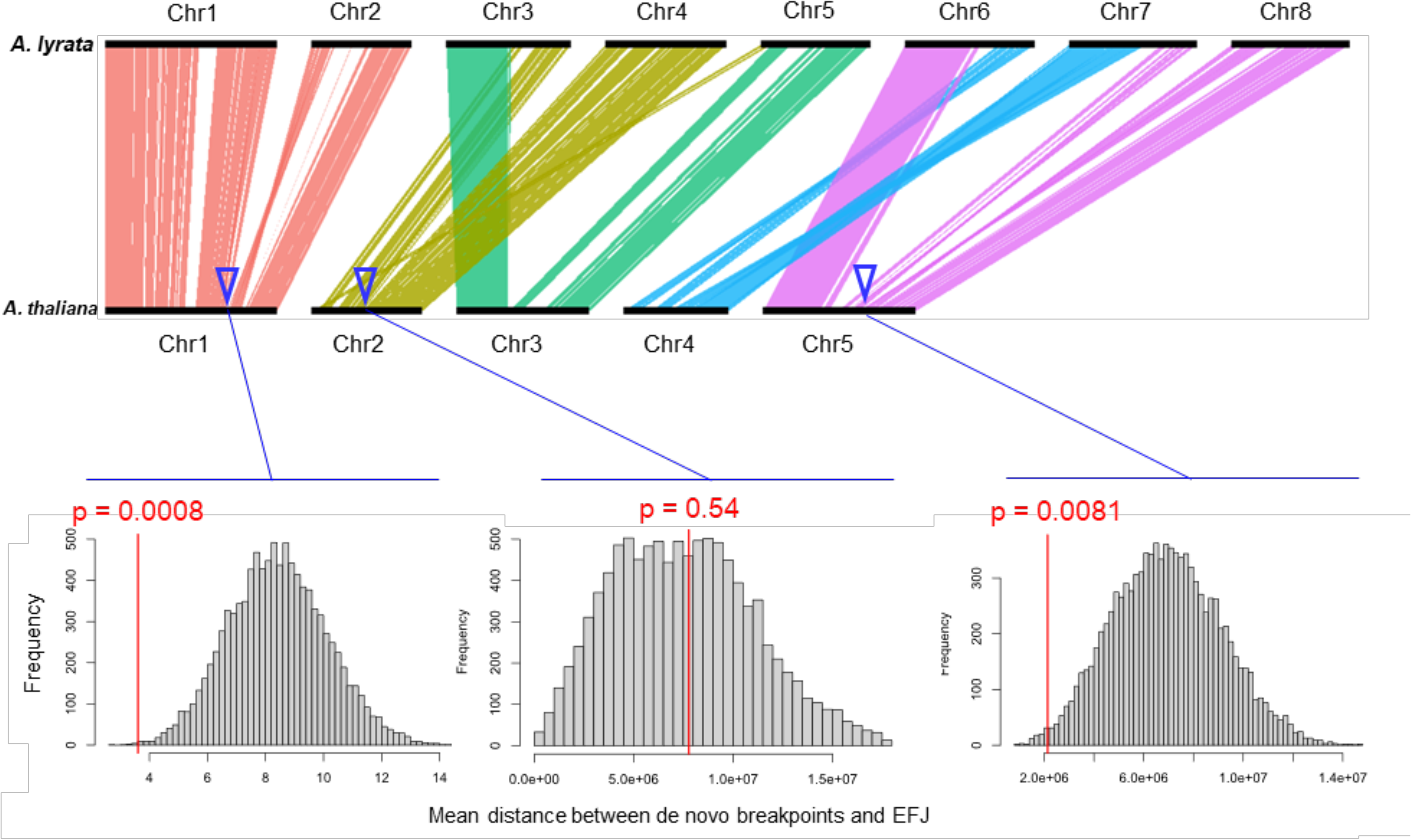
Association between *de novo* breakpoints and ancient end-to-end fusions. Rearrangements that gave rise to A. *thaliana* karyotype are illustrated using synteny between *A. thaliana and A. thaliana* according to ^2^. For each of the three end-to-end fusions that took place in the lineage of *A. thaliana*, we tested whether the detected *de novo* breakpoints are less distant than expected by chance. Grey bars in the plots represent the frequency of the distances between simulated random breakpoints and the ancient fusion breakpoint. The red lines mark the observed distance between the actual *de novo* breakpoints and the ancient fusion breakpoint. The p-values reflect the probability of observing the actual distances in the simulated distribution according to permutation tests.

**Figure S8:**
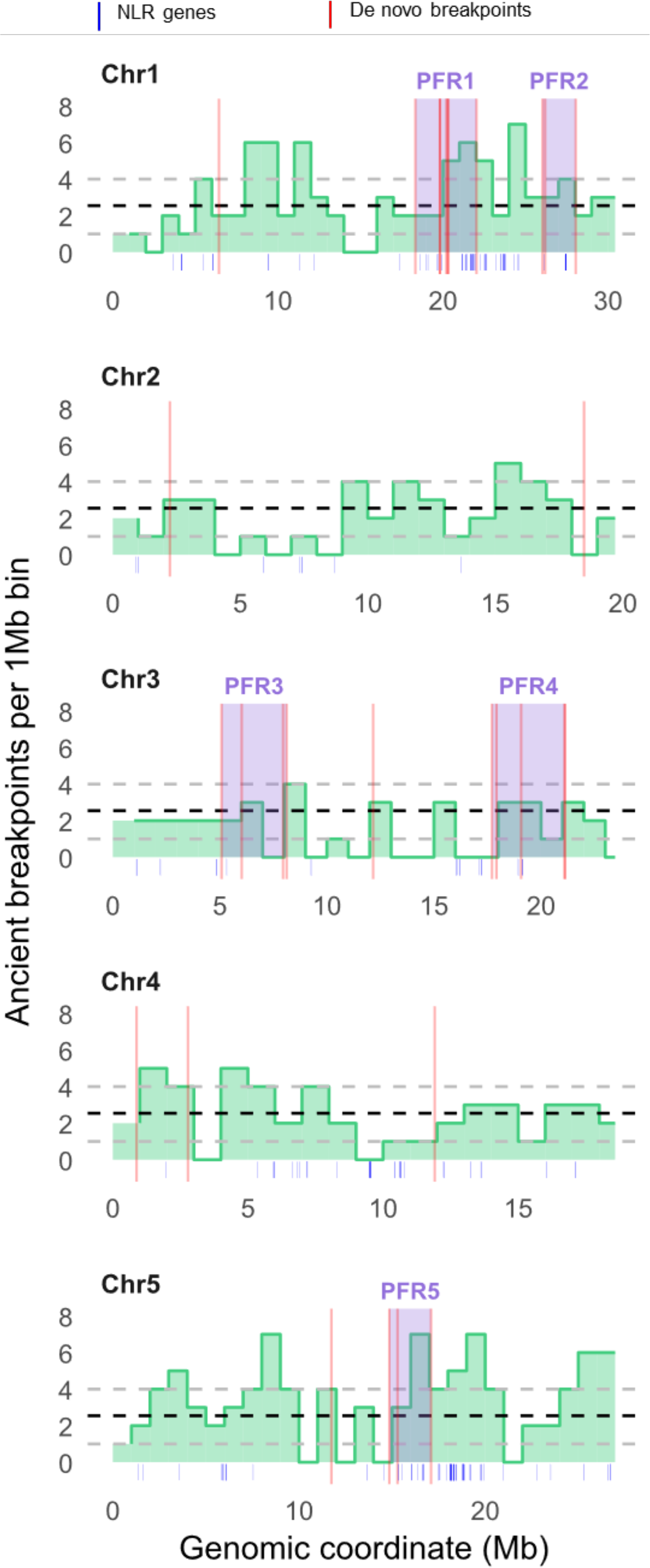
NLR genes and rearrangements (ancient and *de novo*). The position of NLR genes reported in ^3^ (depicted as dark blue lines below the plots) are shown along with a density map of the breakpoints for ancient genome rearrangements identified during 25 MY of Brassicaceae evolution^4^. The genome is divided into bins of 1 Mb. Black dashed lines indicate the genome average of ancient breakpoints per Mb whereas the lower and upper grey dashed lines indicate the first and the fourth quantiles (respectively) for ancient breakpoints per megabase. This density map was made using the TAIR10.1 genome.

**Figure S9:**
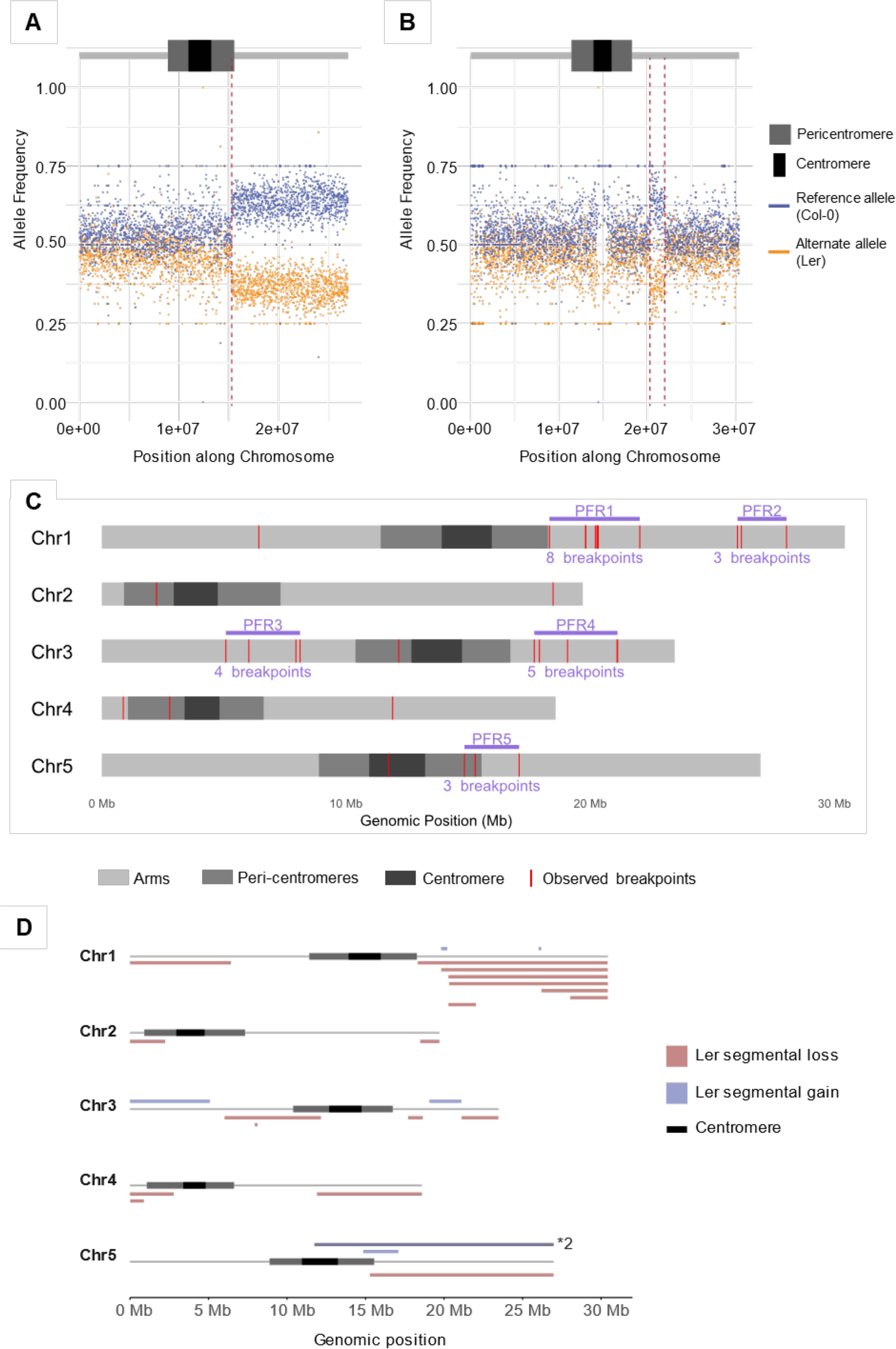
*De novo* breakpoints in segmental aneuploidies as in Figure 2, but in this case mapped to the TAIR10.1 (Col-0) genome. (A,B) Two examples of how breakpoints are mapped using allele frequencies at high resolution. Every data point represents a 10 kb window across the genome. (C) Breakpoint map of the TAIR10.1 genome, including the identified PFRs. (D) Size and position of all identified segmental gains (blue, above chromosome line) and losses (red, below chromosome line) across all chromosomes. *2 indicates a fragment that was gained twice within a single individual.

## Supplemental tables

**Supplemental Table S1:** Summary of counts of euploid and aneuploid gametes (all aneuploidies considered).

| Crosses | Parent of origin | Euploid | Aneuploid | Total contributions |
| --- | --- | --- | --- | --- |
| 1,3 and 5 | Female Col | 90 | 7 | 97 |
| 1,3 and 5 | Male Ler | 110 | 21 | 131 |
| 2,4 and 6 | Female Ler | 101 | 30 | 131 |
| 2,4 and 6 | Male Col | 88 | 9 | 97 |
|  | total | 389 | 67 | 456 |

**Supplemental Table S2:**
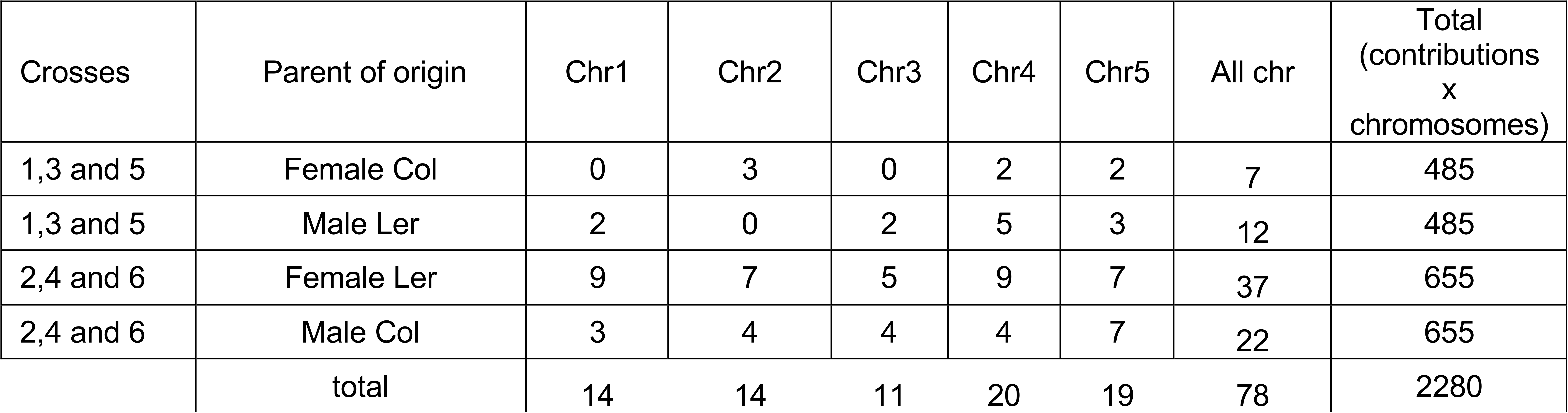
Summary of counts of gametes affected by any kind of aneuploidy for a given chromosome.

**Supplemental Table S3:** Summary of counts of gametes affected by full-length aneuploidy for a given chromosome.

| Crosses | Parent of origin | Chr1 | Chr2 | Chr3 | Chr4 | Chr5 | All chr | Total<br>(contributions<br>x<br>chromosomes) |
| --- | --- | --- | --- | --- | --- | --- | --- | --- |
| 1,3 and 5 | Female Col | 0 | 3 | 0 | 2 | 2 | 7 | 485 |
| 1,3 and 5 | Male Ler | 2 | 0 | 1 | 5 | 3 | 11 | 485 |
| 2,4 and 6 | Female Ler | 1 | 5 | 3 | 6 | 5 | 20 | 655 |
| 2,4 and 6 | Male Col | 3 | 4 | 4 | 4 | 7 | 22 | 655 |
|  | total | 6 | 12 | 8 | 17 | 17 | 60 | 2280 |

**Supplemental Table S4:** Summary of counts of gametes affected by segmental aneuploidy for a given chromosome.

| Crosses | Parent of origin | Chr1 | Chr2 | Chr3 | Chr4 | Chr5 | All chr | Total (contributions x chromosomes) |
| --- | --- | --- | --- | --- | --- | --- | --- | --- |
| 1,3 and 5 | Female Col | 0 | 0 | 0 | 0 | 0 | 0 | 485 |
| 1,3 and 5 | Male Ler | 0 | 0 | 1 | 0 | 1 | 2 | 485 |
| 2,4 and 6 | Female Ler | 8 | 2 | 2 | 3 | 2 | 17 | 655 |
| 2,4 and 6 | Male Col | 0 | 0 | 0 | 0 | 0 | 0 | 655 |
|  | total | 8 | 2 | 3 | 3 | 3 | 19 | 2280 |

**Supplemental Table S5:** Summary of counts of gametes that are affected by full-length gains for a given chromosome.

| Crosses | Parent of origin | Chr1 | Chr2 | Chr3 | Chr4 | Chr5 | All chr | Total (contributions x chromosomes) |
| --- | --- | --- | --- | --- | --- | --- | --- | --- |
| 1,3 and 5 | Female Col | 0 | 1 | 0 | 2 | 0 | 3 | 485 |
| 1,3 and 5 | Male Ler | 0 | 0 | 1 | 5 | 3 | 9 | 485 |
| 2,4 and 6 | Female Ler | 1 | 1 | 2 | 4 | 3 | 11 | 655 |
| 2,4 and 6 | Male Col | 2 | 4 | 0 | 4 | 7 | 17 | 655 |
|  | total | 3 | 6 | 3 | 15 | 13 | 40 | 2280 |

**Supplemental Table S6:** Summary of counts of gametes that are affected by full-length losses for a given chromosome.

| Crosses | Parent of origin | Chr1 | Chr2 | Chr3 | Chr4 | Chr5 | All chr | Total (contributions x chromosomes) |
| --- | --- | --- | --- | --- | --- | --- | --- | --- |
| 1,3 and 5 | Female Col | 0 | 2 | 0 | 0 | 2 | 4 | 485 |
| 1,3 and 5 | Male Ler | 2 | 0 | 0 | 0 | 0 | 2 | 485 |
| 2,4 and 6 | Female Ler | 0 | 4 | 1 | 2 | 2 | 9 | 655 |
| 2,4 and 6 | Male Col | 1 | 0 | 4 | 0 | 0 | 5 | 655 |
|  | total | 3 | 6 | 5 | 2 | 4 | 20 | 2280 |

**Supplemental Table S7:** Summary of gametes that are affected by segmental gains for a given chromosome.

| Crosses | Parent of origin | Chr1 | Chr2 | Chr3 | Chr4 | Chr5 | All chr | Total (contributions x chromosomes) |
| --- | --- | --- | --- | --- | --- | --- | --- | --- |
| 1,3 and 5 | Female Col | 0 | 0 | 0 | 0 | 0 | 0 | 485 |
| 1,3 and 5 | Male Ler | 0 | 0 | 0 | 0 | 1 | 0 | 485 |
| 2,4 and 6 | Female Ler | 2 | 0 | 2 | 0 | 1 | 5 | 655 |
| 2,4 and 6 | Male Col | 2 | 0 | 1 | 0 | 0 | 3 | 655 |
|  | total | 4 | 0 | 3 | 0 | 2 | 8 | 2280 |

**Supplemental Table S8:** Summary of gametes that are affected by segmental losses for a given chromosome.

| Crosses | Parent of origin | Chr1 | Chr2 | Chr3 | Chr4 | Chr5 | All chr | Total (contributions x chromosomes) |
| --- | --- | --- | --- | --- | --- | --- | --- | --- |
| 1,3 and 5 | Female Col | 0 | 0 | 0 | 0 | 0 | 0 | 485 |
| 1,3 and 5 | Male Ler | 0 | 0 | 1 | 0 | 0 | 1 | 485 |
| 2,4 and 6 | Female Ler | 8 | 2 | 1 | 3 | 1 | 15 | 655 |
| 2,4 and 6 | Male Col | 0 | 0 | 0 | 0 | 0 | 0 | 655 |
|  | total | 8 | 2 | 2 | 3 | 1 | 16 | 2280 |

**Supplemental Table S9:** Segmental breakpoints mapped onto Ler chromosomes.

| Chr | Break | Sample |
| --- | --- | --- |
| Chr1 | 6480000 | P6_10D |
| Chr1 | 17350001 | P6_26C |
| Chr1 | 18800001 | P6_23D |
| Chr1 | 18800001 | P6_27C |
| Chr1 | 19220001 | P6_27C |
| Chr1 | 19290001 | P4_27B |
| Chr1 | 19310001 | P6_16B |
| Chr1 | 20960000 | P6_16B |
| Chr1 | 24970001 | P4_32C |
| Chr1 | 25130001 | P4_32C |
| Chr1 | 26990001 | P4_17A |
| Chr2 | 2260000 | P6_13B |
| Chr2 | 17820001 | P6_16B |
| Chr3 | 530001 | P6_16B |
| Chr3 | 5390000 | P6_16B |
| Chr3 | 6080001 | P3_1B |
| Chr3 | 7940001 | P6_13B |
| Chr3 | 8320000 | P6_13B |
| Chr3 | 12350000 | P3_1B |
| Chr3 | 16900001 | P6_13B |
| Chr3 | 17800000 | P6_13B |
| Chr3 | 18570001 | P6_13B |
| Chr3 | 20320000 | P6_13B |
| Chr4 | 880000 | P4_24C |
| Chr4 | 2580000 | P6_13B |
| Chr4 | 11270001 | P2_1A |
| Chr5 | 11720001 | P4_12D |
| Chr5 | 14200001 | P3_14D |
| Chr5 | 14600001 | P4_24C |
| Chr5 | 16500000 | P3_14D |

**Supplemental Table S10:** Identified losses and gains in chromosomes affected by segmental aneuploidy.

| Chr | start | end | length | sample | type |
| --- | --- | --- | --- | --- | --- |
| Chr4 | 11270001 | 17980856 | 6710855 | P2_1A | Loss |
| Chr5 | 14200001 | 16500000 | 2299999 | P3_14D | Gain |
| Chr3 | 6080001 | 12350000 | 6269999 | P3_1B | Loss |
| Chr5 | 11720001 | 26372590 | 14652589 | P4_12D | Gain (double) |
| Chr1 | 26990001 | 29387870 | 2397869 | P4_17A | Loss |
| Chr4 | 1 | 880000 | 879999 | P4_24C | Loss |
| Chr5 | 14600001 | 26372590 | 11772589 | P4_24C | Loss |
| Chr1 | 19290001 | 29387870 | 10097869 | P4_27B | Loss |
| Chr1 | 24970001 | 25130000 | 159999 | P4_32C | Gain |
| Chr1 | 25130001 | 29387870 | 4257869 | P4_32C | Loss |
| Chr1 | 1 | 6480000 | 6479999 | P6_10D | Loss |
| Chr2 | 1 | 2260000 | 2259999 | P6_13B | Loss |
| Chr3 | 7940001 | 8320000 | 379999 | P6_13B | Loss |
| Chr3 | 16900001 | 17800000 | 899999 | P6_13B | Loss |
| Chr3 | 18570001 | 20320000 | 1749999 | P6_13B | Gain |
| Chr3 | 20320001 | 22588203 | 2268202 | P6_13B | Loss |
| Chr4 | 1 | 2580000 | 2579999 | P6_13B | Loss |
| Chr1 | 19310001 | 20960000 | 1649999 | P6_16B | Loss |
| Chr2 | 17820001 | 19037554 | 1217553 | P6_16B | Loss |
| Chr3 | 530001 | 5390000 | 4859999 | P6_16B | Gain |
| Chr1 | 18800001 | 29387870 | 10587869 | P6_23D | Loss |
| Chr1 | 17350001 | 29387870 | 12037869 | P6_26C | Loss |
| Chr1 | 18800001 | 19220000 | 419999 | P6_27C | Gain |
| Chr1 | 19220001 | 29387870 | 10167869 | P6_27C | Loss |

**Supplemental Table S11:** Location of identified PFRs in Ler reference.

| PFR | start | end | length | number of breakpoints |
| --- | --- | --- | --- | --- |
| PFR1 | 17350001 | 20960000 | 3609999 | 7 |
| PFR2 | 24970001 | 26990001 | 2020000 | 4 |
| PFR3 | 5390000 | 8320000 | 2930000 | 4 |
| PFR4 | 16900001 | 20320000 | 3419999 | 4 |
| PFR5 | 14200001 | 16500000 | 2299999 | 3 |
| PFR1_core | 18800001 | 19310001 | 510000 | 5 |

**Supplemental Table S12:** End-to-end fusions between *A. lyrata* chromosomes (Aly_Chrs) to form the *A. thaliana* chromosomes (Ath_ Chrs) inferred by synteny between *A. thaliana* and *A. lyrata* according to ^2^. The fusion breakpoint coordinate in TAIR (TAIR_coordinate) was established as the intermediate position between the southernmost and the northernmost homology synteny blocks from ^2^ of the two *A. lyrata* chromosomes involved in the fusion. To transfer this coordinate to Ler, we used the proximal gene to the breakpoint established in TAIR (TAIR_gene), found the corresponding synthenic ortholog in Ler (LerZ gene) and used its end coordinate (LerZ_coordinate).

| Aly_Chrs | Ath_Chrs | TAIR_coordinate | TAIR_gene | LerZ_gene | LerZ_coordinate |
| --- | --- | --- | --- | --- | --- |
| Chr1_Chr2 | Chr1 | 21431160 | AT1G57870 | AT1G51900 | 20439472 |
| Chr3_Chr4 | Chr2 | 8995763 | AT2G20900 | AT2G20920 | 8312480 |
| Chr7_Chr8 | Chr5 | 16833899 | AT5G42120 | AT5G39930 | 16282231 |

**Supplemental Table S13:** Summary of the analysis of rearrangemebts betwen macrosynteny blocks during the evolutionary history of Brasicaceae. We used the blocks and the rearrangements identified by Liu et al 2024. Each block is delimited by the first and last syntenic gene (ID_start and ID_end, respectively). We used the start (start_bp) and end (end_bp) coordinates of the first and last syntenic genes in Ler, respectively, as the boundaries of each block. For each block, we registered the chromosome (Chr) where it is located and its size in Mb (size_mb). For each block boundary (north and south) we counted the number of times that the adjacent block changed due to all of the rearrangements registered by ^4^ (north swaps and south swaps, respectively). This information was used to make Figure 3C.

| block | ID_start | start_bp | ID_end | end_bp | Chr | Size (mb) | North swaps | South swaps |
| --- | --- | --- | --- | --- | --- | --- | --- | --- |
| A1 | AT1G01020 | 7313 | AT1G08100 | 2533125 | Chr1 | 2.525812 | 1 | 1 |
| A2 | AT1G08110 | 2539320 | AT1G12970 | 4455757 | Chr1 | 1.916437 | 1 | 1 |
| A3 | AT1G12970 | 4453503 | AT1G16610 | 5704433 | Chr1 | 1.25093 | 1 | 2 |
| A4 | AT1G16630 | 5704983 | AT1G19840 | 6880587 | Chr1 | 1.175604 | 2 | 2 |
| B1 | AT1G19850 | 6893982 | AT1G24260 | 8577010 | Chr1 | 1.683028 | 2 | 2 |
| B2 | AT1G24260 | 8574676 | AT1G24260 | 8577010 | Chr1 | 0.002334 | 4 | 2 |
| B3 | AT1G27290 | 9396005 | AT1G30755 | 10852366 | Chr1 | 1.456361 | 4 | 2 |
| B4 | AT1G30760 | 10861531 | AT1G32750 | 11792228 | Chr1 | 0.930697 | 2 | 4 |
| B5 | AT1G32760 | 11792993 | AT1G37130 | 14113481 | Chr1 | 2.320488 | 3 | 2 |
| C1 | AT1G43020 | 15186034 | AT1G47920 | 16700357 | Chr1 | 1.514323 | 3 | 2 |
| C2 | AT1G47960 | 16707630 | AT1G53710 | 19046291 | Chr1 | 2.338661 | 2 | 2 |
| C3 | AT1G53720 | 19046466 | AT1G56190 | 20006318 | Chr1 | 0.959852 | 2 | 3 |
| D1 | AT1G56210 | 20011291 | AT1G61210 | 21487664 | Chr1 | 1.476373 | 6 | 3 |
| D2 | AT1G61215 | 21488295 | AT1G64670 | 22975250 | Chr1 | 1.486955 | 2 | 2 |
| E1 | AT1G64960 | 23071825 | AT1G67270 | 24110829 | Chr1 | 1.039004 | 4 | 3 |
| E2 | AT1G67280 | 24113306 | AT1G71100 | 25748415 | Chr1 | 1.635109 | 3 | 3 |
| E3 | AT1G71110 | 25750714 | AT1G78310 | 28438216 | Chr1 | 2.687502 | 4 | 2 |
| E4 | AT1G78320 | 28438116 | AT1G79720 | 28966997 | Chr1 | 0.528881 | 2 | 1 |
| E5 | AT1G79730 | 28968473 | AT1G80950 | 29382002 | Chr1 | 0.413529 | 1 | 2 |
| KL1 | AT2G01060 | 5279 | AT2G05160 | 1845851 | Chr2 | 1.840572 | 2 | 1 |
| G | AT2G05170 | 1853956 | AT2G07690 | 162711 | Chr2 | -1.69125 | 3 | 3 |
| H | AT2G10940 | 3767203 | AT2G20900 | 8312480 | Chr2 | 4.545277 | 1 | 1 |
| I1 | AT2G20930 | 8319159 | AT2G25260 | 10078428 | Chr2 | 1.759269 | 4 | 2 |
| I2 | AT2G25270 | 10080273 | AT2G27550 | 11084971 | Chr2 | 1.004698 | 2 | 2 |
| I3 | AT2G27550 | 11083527 | AT2G31050 | 12531968 | Chr2 | 1.448441 | 2 | 1 |
| J1 | AT2G31040 | 12527996 | AT2G35850 | 14388873 | Chr2 | 1.860877 | 1 | 2 |
| J2 | AT2G35860 | 14391359 | AT2G37660 | 15127667 | Chr2 | 0.736308 | 4 | 1 |
| J3 | AT2G37670 | 15127790 | AT2G41410 | 16591113 | Chr2 | 1.463323 | 1 | 3 |
| J4 | AT2G41430 | 16597538 | AT2G48150 | 19028914 | Chr2 | 2.431376 | 3 | 2 |
| F1 | AT3G01015 | 1756 | AT3G07530 | 2424057 | Chr3 | 2.422301 | 2 | 2 |
| F2 | AT3G07540 | 2424109 | AT3G12180 | 3921285 | Chr3 | 1.497176 | 2 | 2 |
| F3 | AT3G12190 | 3922350 | AT3G16010 | 5480603 | Chr3 | 1.558253 | 2 | 2 |
| F4 | AT3G16030 | 5483288 | AT3G25520 | 9327771 | Chr3 | 3.844483 | 3 | 4 |
| KL2 | AT3G25540 | 9330836 | AT3G32960 | 22581 | Chr3 | -9.30826 | 1 | 3 |
| MN1 | AT3G42180 | 12595179 | AT3G52970 | 18830319 | Chr3 | 6.23514 | 3 | 3 |
| MN2 | AT3G52980 | 18831814 | AT3G56550 | 20171188 | Chr3 | 1.339374 | 3 | 1 |
| MN3 | AT3G56570 | 20176246 | AT3G59550 | 21211338 | Chr3 | 1.035092 | 1 | 2 |
| MN4 | AT3G59570 | 21211558 | AT3G63530 | 22572163 | Chr3 | 1.360605 | 2 | 0 |
| O1 | AT4G00026 | 15507 | AT4G03190 | 1453087 | Chr4 | 1.43758 | 2 | 4 |
| O2 | AT4G03200 | 1453552 | AT4G05450 | 1717043 | Chr4 | 0.263491 | 1 | 4 |
| P1 | AT4G08170 | 4470704 | AT4G12620 | 6822203 | Chr4 | 2.351499 | 5 | 4 |
| P2 | AT4G09680 | 5374974 | AT4G12620 | 6822203 | Chr4 | 1.447229 | 2 | 1 |
| T | AT4G12700 | 6842329 | AT4G16250 | 8628764 | Chr4 | 1.786435 | 3 | 2 |
| U1 | AT4G16250 | 8624375 | AT4G24150 | 11891372 | Chr4 | 3.266997 | 1 | 1 |
| U2 | AT4G24160 | 11891546 | AT4G27730 | 13177479 | Chr4 | 1.285933 | 2 | 3 |
| U3 | AT4G27740 | 13180024 | AT4G33010 | 15286997 | Chr4 | 2.106973 | 3 | 1 |
| U4 | AT4G33000 | 15280302 | AT4G35730 | 16309593 | Chr4 | 1.029291 | 1 | 2 |
| U5 | AT4G35733 | 16310520 | AT4G38100 | 17278479 | Chr4 | 0.967959 | 2 | 1 |
| U6 | AT4G38120 | 17278737 | AT4G40090 | 17977392 | Chr4 | 0.698655 | 1 | 1 |
| R1 | AT5G01010 | 1517 | AT5G06730 | 2099748 | Chr5 | 2.098231 | 1 | 2 |
| R2 | AT5G06740 | 2100763 | AT5G08535 | 2782828 | Chr5 | 0.682065 | 2 | 2 |
| R3 | AT5G08540 | 2783234 | AT5G13380 | 4335420 | Chr5 | 1.552186 | 2 | 3 |
| R4 | AT5G13390 | 4335738 | AT5G19340 | 6505963 | Chr5 | 2.170225 | 3 | 2 |
| R5 | AT5G19350 | 6508578 | AT5G23000 | 7701553 | Chr5 | 1.192975 | 3 | 4 |
| Q1 | AT5G23010 | 7705999 | AT5G26230 | 9182509 | Chr5 | 1.47651 | 2 | 5 |
| Q2 | AT5G26230 | 9181246 | AT5G30510 | 11597923 | Chr5 | 2.416677 | 4 | 4 |
| S1 | AT5G32470 | 11748000 | AT5G39880 | 15405215 | Chr5 | 3.657215 | 3 | 3 |
| S2 | AT5G39890 | 15408228 | AT5G42110 | 16279381 | Chr5 | 0.871153 | 4 | 3 |
| V | AT5G42130 | 16282332 | AT5G47810 | 18756614 | Chr5 | 2.474282 | 4 | 5 |
| W1 | AT5G47820 | 18759139 | AT5G49610 | 19538952 | Chr5 | 0.779813 | 6 | 1 |
| W2 | AT5G49620 | 19543512 | AT5G56540 | 22272034 | Chr5 | 2.728522 | 4 | 2 |
| W3 | AT5G56550 | 22273955 | AT5G60800 | 23845016 | Chr5 | 1.571061 | 2 | 3 |
| X1 | AT5G60805 | 23847545 | AT5G63090 | 24698528 | Chr5 | 0.850983 | 1 | 3 |
| X2 | AT5G63100 | 24701855 | AT5G65925 | 25763253 | Chr5 | 1.061398 | 3 | 2 |
| X3 | AT5G65930 | 25763567 | AT5G67640 | 26366528 | Chr5 | 0.602961 | 2 | 2 |

**Supplemental Table S14:**
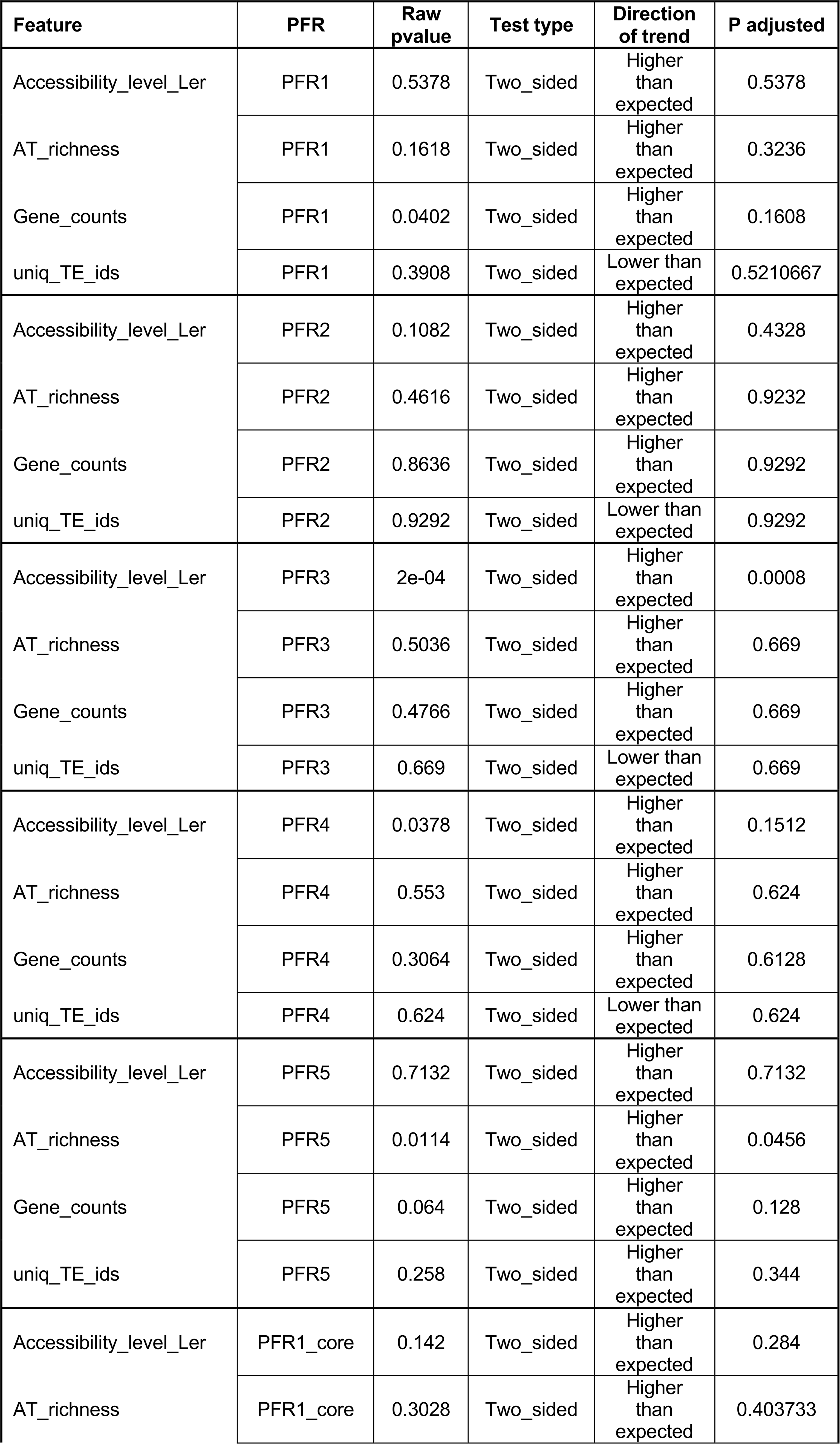
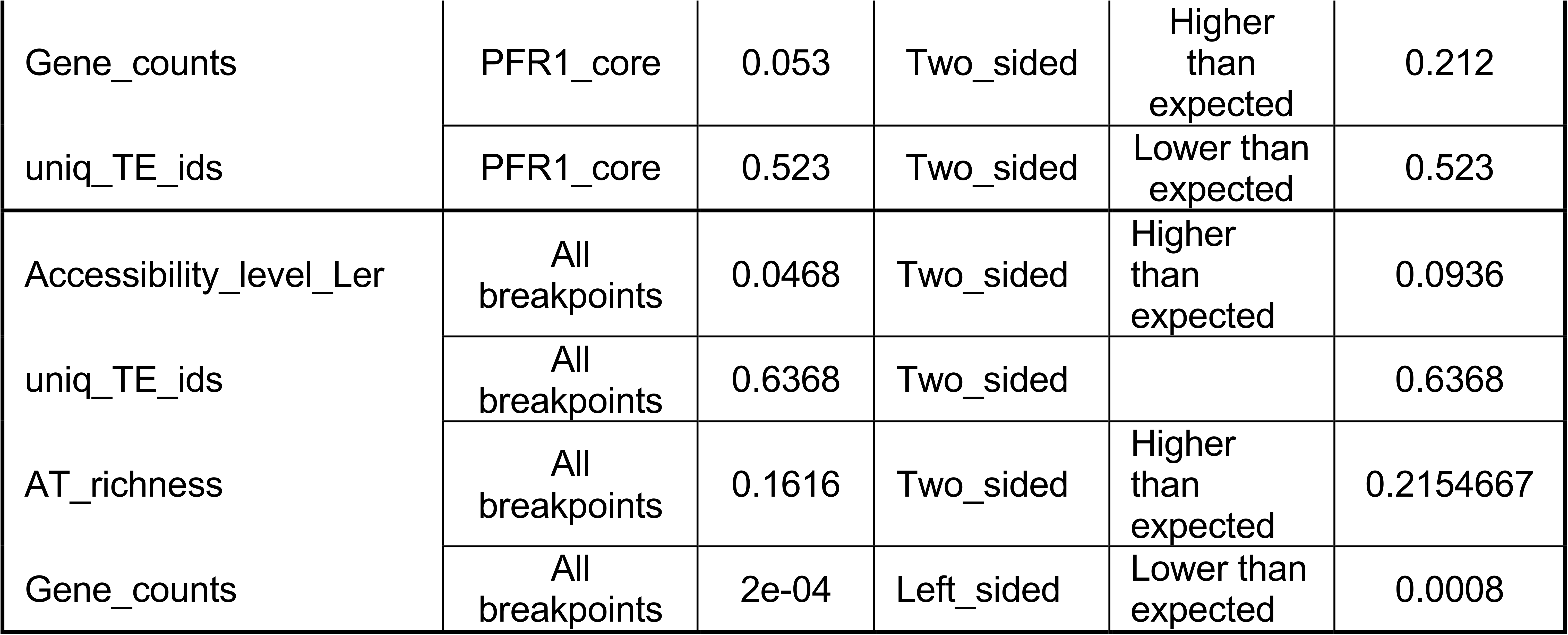
Summary of the results of analyses of genomic features of PFRs.

| Feature | PFR | Raw pvalue | Test type | Direction of trend | P adjusted |
| --- | --- | --- | --- | --- | --- |
| Accessibility_level_Ler | PFR1 | 0.5378 | Two_sided | Higher than expected | 0.5378 |
| AT_richness | PFR1 | 0.1618 | Two_sided | Higher than expected | 0.3236 |
| Gene_counts | PFR1 | 0.0402 | Two_sided | Higher than expected | 0.1608 |
| uniq_TE_ids | PFR1 | 0.3908 | Two_sided | Lower than expected | 0.5210667 |
| Accessibility_level_Ler | PFR2 | 0.1082 | Two_sided | Higher than expected | 0.4328 |
| AT_richness | PFR2 | 0.4616 | Two_sided | Higher than expected | 0.9232 |
| Gene_counts | PFR2 | 0.8636 | Two_sided | Higher than expected | 0.9292 |
| uniq_TE_ids | PFR2 | 0.9292 | Two_sided | Lower than expected | 0.9292 |
| Accessibility_level_Ler | PFR3 | 2e-04 | Two_sided | Higher than expected | 0.0008 |
| AT_richness | PFR3 | 0.5036 | Two_sided | Higher than expected | 0.669 |
| Gene_counts | PFR3 | 0.4766 | Two_sided | Higher than expected | 0.669 |
| uniq_TE_ids | PFR3 | 0.669 | Two_sided | Lower than expected | 0.669 |
| Accessibility_level_Ler | PFR4 | 0.0378 | Two_sided | Higher than expected | 0.1512 |
| AT_richness | PFR4 | 0.553 | Two_sided | Higher than expected | 0.624 |
| Gene_counts | PFR4 | 0.3064 | Two_sided | Higher than expected | 0.6128 |
| uniq_TE_ids | PFR4 | 0.624 | Two_sided | Lower than expected | 0.624 |
| Accessibility_level_Ler | PFR5 | 0.7132 | Two_sided | Higher than expected | 0.7132 |
| AT_richness | PFR5 | 0.0114 | Two_sided | Higher than expected | 0.0456 |
| Gene_counts | PFR5 | 0.064 | Two_sided | Higher than expected | 0.128 |
| uniq_TE_ids | PFR5 | 0.258 | Two_sided | Higher than expected | 0.344 |
| Accessibility_level_Ler | PFR1_core | 0.142 | Two_sided | Higher than expected | 0.284 |
| AT_richness | PFR1_core | 0.3028 | Two_sided | Higher than expected | 0.403733 |
| Gene_counts | PFR1_core | 0.053 | Two_sided | Higher than expected | 0.212 |
| uniq_TE_ids | PFR1_core | 0.523 | Two_sided | Lower than expected | 0.523 |
| Accessibility_level_Ler | All breakpoints | 0.0468 | Two_sided | Higher than expected | 0.0936 |
| uniq_TE_ids | All breakpoints | 0.6368 | Two_sided |  | 0.6368 |
| AT_richness | All breakpoints | 0.1616 | Two_sided | Higher than expected | 0.2154667 |
| Gene_counts | All breakpoints | 2e-04 | Left_sided | Lower than expected | 0.0008 |

**Supplemental Table S15:**
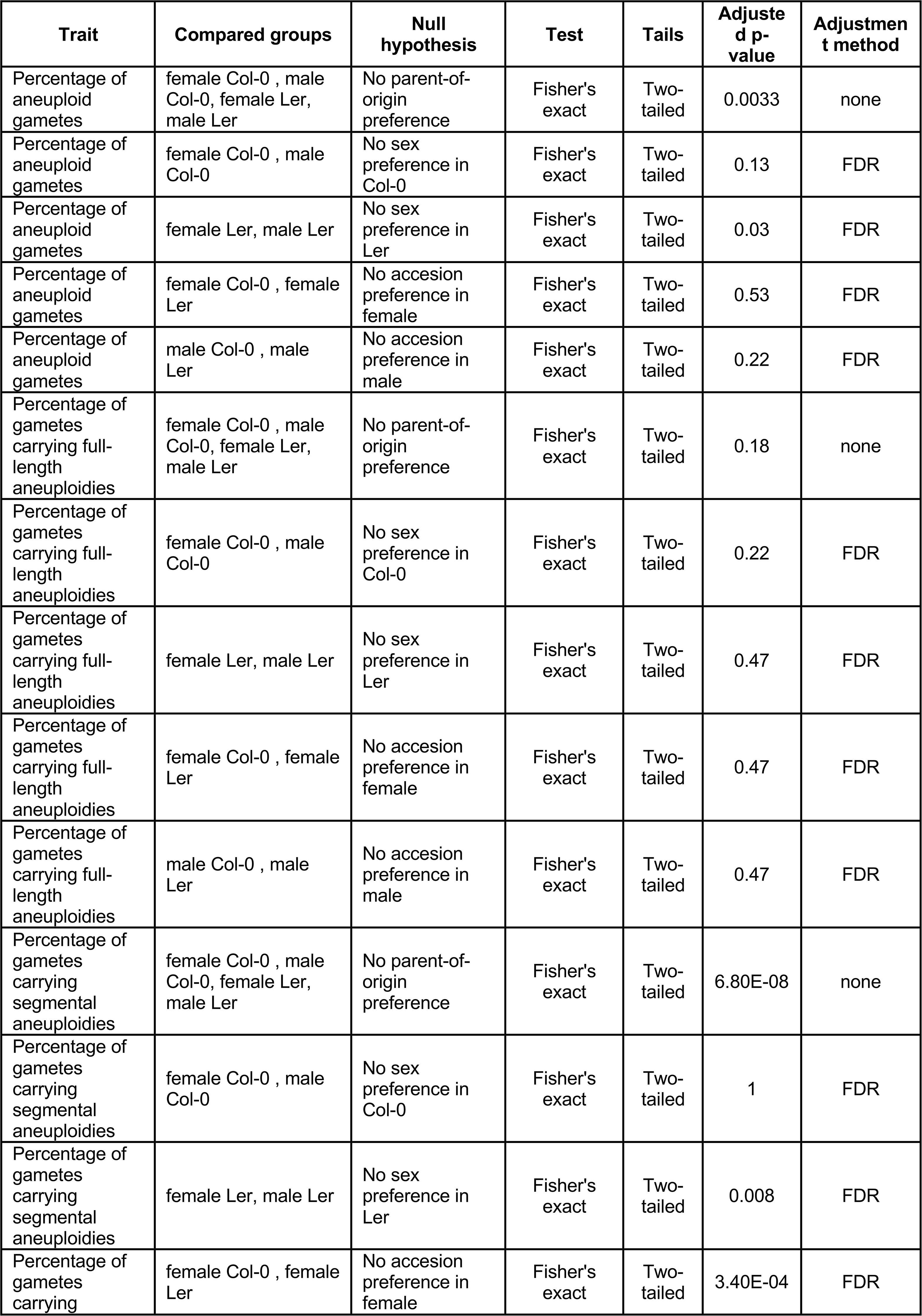
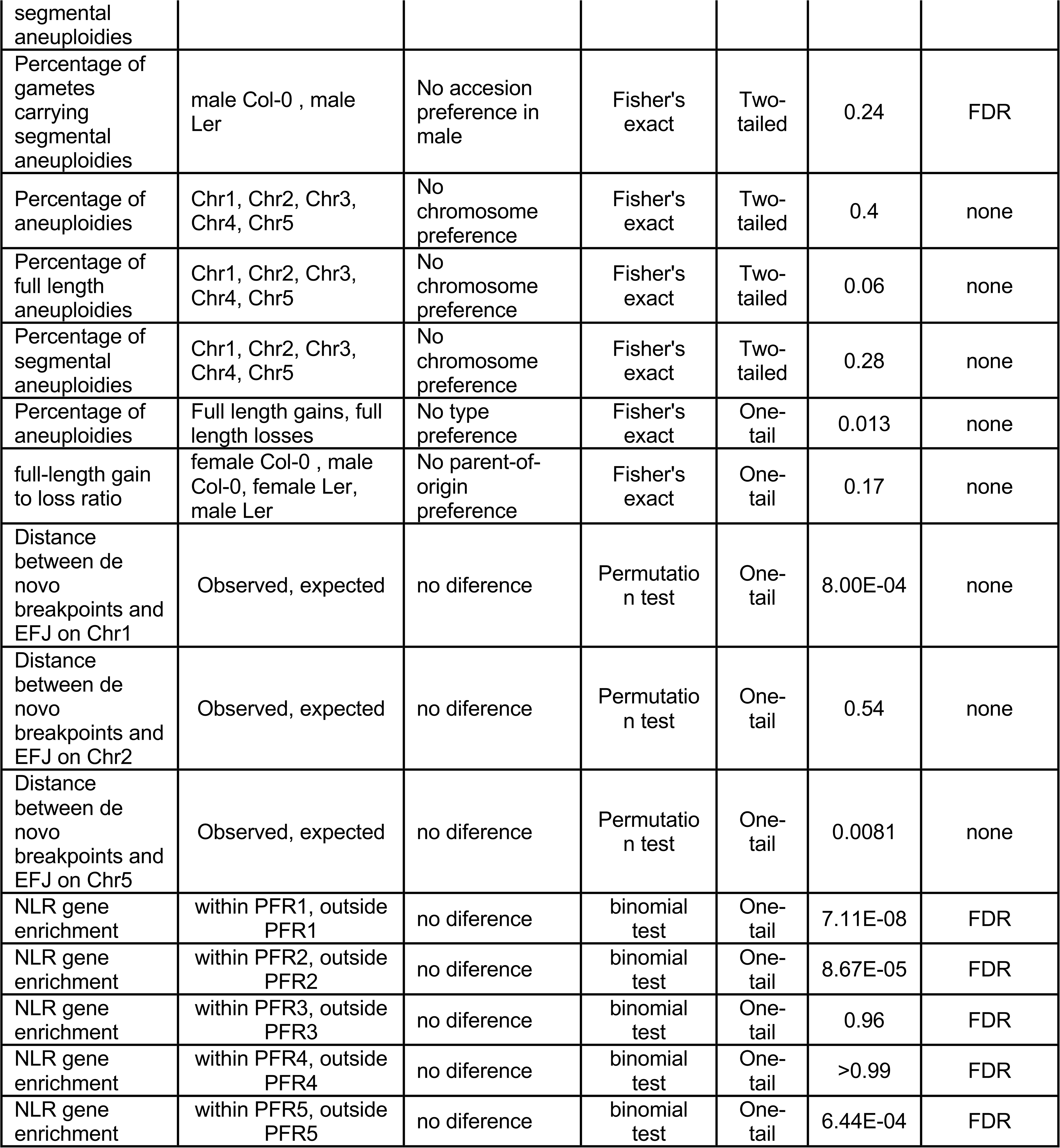
Summary of the statitical analysis (excluding GO and genomic features). When appropriate false discovery rate (FDR) p-value adjustment method was used.

